# Feasibility analysis of semiconductor voltage nanosensors for neuronal membrane potential sensing

**DOI:** 10.1101/838342

**Authors:** Anastasia Ludwig, Pablo Serna, Lion Morgenstein, Gaoling Yang, Omri Bar-Elli, Gloria Ortiz, Evan Miller, Dan Oron, Asaf Grupi, Shimon Weiss, Antoine Triller

## Abstract

In the last decade, optical imaging methods have significantly improved our understanding of the information processing principles in the brain. Although many promising tools have been designed, sensors of membrane potential are lagging behind the rest. Semiconductor nanoparticles are an attractive alternative to classical voltage indicators, such as voltage-sensitive dyes and proteins. Such nanoparticles exhibit high sensitivity to external electric fields via the quantum-confined Stark effect. Here we report the development of lipid-coated semiconductor voltage-sensitive nanorods (vsNRs) that self-insert into the neuronal membrane. We describe a workflow to detect and process the photoluminescent signal of vsNRs after wide-field time-lapse recordings. We also present data indicating that vsNRs are feasible for sensing membrane potential in neurons at a single-particle level. This shows the potential of vsNRs for detection of neuronal activity with unprecedentedly high spatial and temporal resolution.

## Introduction

Electrochemical communication between neurons lies in the basis of brain function. Registering the electrical activity of numerous neurons embedded in a neural circuit is a prerequisite for understanding how their activity generates behavior. Classical electrophysiological approaches do not allow probing for a large neuronal assembly, whereas electric or magnetic encephalograms provide wider spatial coverage, but poor spatial resolution. ^1^ Optical imaging offers an alternative approach, namely visualization of neuronal activity using voltage- and calcium-sensitive fluorescent reporters.^2^ In the last decade, the rapid development of calcium indicators and their wide-spread application has revolutionized the field. ^3^ However, oscillations of intracellular calcium concentration are an indirect measure of electrical activity, with limited accuracy due to slow dynamics and interference of intracellular signaling machinery. Thus, the development of competent membrane potential indicators remains highly desirable.

An ideal voltage sensor should provide a combination of high spatial resolution with sub-millisecond kinetics. Much effort has been made to develop voltage-sensitive organic dyes,^4–8^ reviewed in,^9–11^ genetically-encoded voltage-sensitive proteins,^12–18^ reviewed in,^2,19–21^ as well as hybrid sensors that combine advantages of dyes and proteins.^22–24^ The latest generation of voltage sensors yielded successful physiological experiments *in vivo*.^16,25–31^ Despite substantial progress, numerous problems, such as low brightness, poor photostability, and slow kinetics, are still to be overcome. ^32–34^

Semiconductor nanoparticles offer an alternative to classical voltage sensors.^35–38^ They are highly photoluminescent and capable of sensing electric field through the quantum-confined Stark effect (QCSE).^39^ QCSE originates from a separation of confined photoexcited charges creating a dipole opposing the external electric field that results in a spectral shift as well as in changes of the quantum yield and photoluminescence (PL) lifetime of the particle. QCSE of symmetric nanoparticles, known as quantum dots (QDs), exhibit quadratic dependence on the field strength.^40–42^ In contrast, asymmetric type-II nanorods (NRs), which consist of a semiconductor shell grown on a QD core, ^43^ exhibit a linear QCSE^44–46^ with higher voltage sensitivity as compared to symmetric QDs. ^37^ If inserted in the plasma membrane orthogonally to the membrane plane, such nanorods could report membrane potential fluctuations via the spectral shift and PL intensity changes. For voltage sensing in neurons, asymmetric type-II NRs are preferable to QDs, since a linear dependence of QCSE on the electric field provides a larger dynamic range for typical activity-dependent fluctuations of the membrane potential between −70 mV and +20 mV.^37^

Voltage nanosensors are potentially more promising than voltage-sensitive dyes and proteins. Apart from higher voltage sensitivity, semiconductor nanoparticles offer superior brightness and resistance to photobleaching in comparison with classical voltage indicators. The exceptional brightness of nanoparticles offers a possibility of single-particle emission measurements. In combination with its nanoscale size, it opens up an avenue for recording of membrane potential from subcellular locations, such as axons or dendritic spines, which remains highly challenging and often unreachable by existing techniques *in vivo*.^47–53^

We have recently demonstrated that NRs are feasible for voltage sensing on a single-particle level at millisecond timescale^54,55^ and can be functionalized with rationally-designed *α*-helical peptides so that they self-insert into a membrane of human embryonic kidney 293 (HEK293) cells and detect evoked voltage fluctuations.^56^ In this work, we present differently functionalized lipid-coated voltage-sensitive NRs (vsNRs) and report a detailed automated workflow that we developed to record vsNR signal from cultured neurons (Fig. 1).

**Figure 1:**
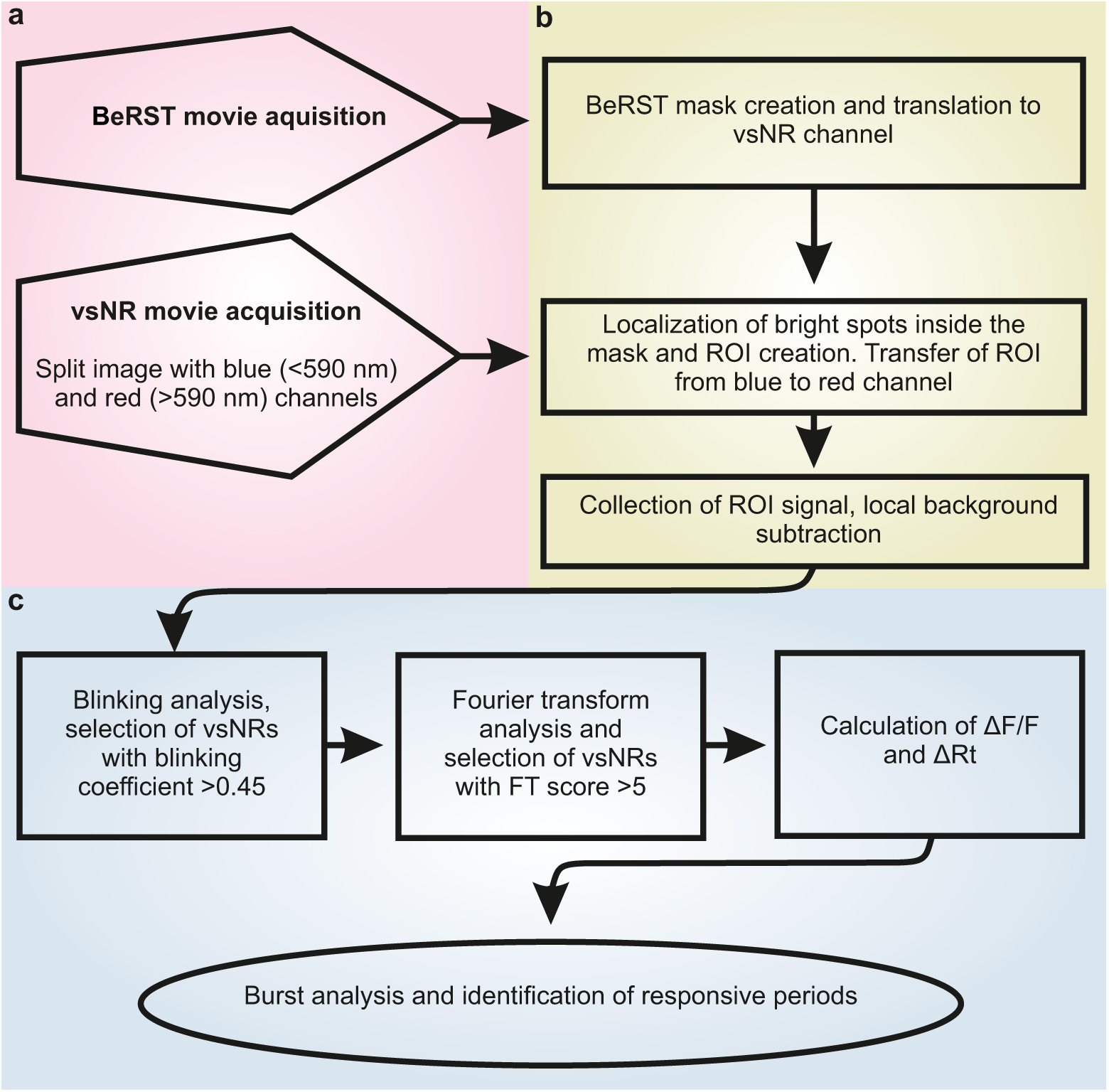
Workflow for vsNR signal recording and analysis. (a) Acquisition of time-lapse movies. (b) Plotting of regions of interest (ROIs) and signal extraction. (c) Signal analysis.

The starting point of our protocol is a movie obtained from a neuronal culture loaded with vsNRs and voltage-sensitive dye BeRST,^57^ where only one cell out of several in the field of view is stimulated through an attached patch electrode to modulate membrane voltage. To identify pixels in the field that belong to the membrane of the stimulated cell, we create a mask using voltage-modulated BeRST fluorescence. Then we identify the location of multiple vsNRs inside the mask, collect their PL signal, as well as measure and subtract local background. The PL signal of a single vsNR contains characteristic temporal intermittency (blinking).^58^ After extracting vsNR signal, we identify single vsNRs based on their blinking pattern and excise “dark states” from the signal.

At the current stage of the technique development, functionalization of NRs with either peptides^56^ or lipids (this study) provides far below 100% membrane insertion. There are no reliable methods to measure insertion in the plasma membrane and the degree of insertion remains uncertain even if the particle stays attached to the cell surface. Thus, most of vsNRs remain unresponsive to voltage modulation during a part or the entire time-trace. At the last stage of analysis, we select responsive vsNRs based on Fourier transform of their signal and identify intervals of time when they were sensitive to voltage modulation.

Overall, we demonstrate that lipid-coated nanorods effectively reach the surface of cultured neurons and adhere to it. Although most of adhered particles remained unresponsive, probably due to membrane insertion failure, a few responsive vsNRs were capable of reporting 60 mV membrane potential fluctuations at a single particle level. We recorded both voltage-related quantum yield changes and maximum emission wavelength shifts.

## Results and Discussion

### vsNR synthesis and functionalization

In this work, we used ZnSe/CdS type-II seeded NRs consisting of a spherical ZnSe core overcoated by a CdS rod (Fig. 2a), synthesized following a modification of a previously reported approach.^59^ According to the transmission electron microscopy (TEM) image (Fig. 2b), NRs have an average length of 16 nm and width of 8 nm. The absorption and emission spectra of this type-II NRs are shown in Fig. 2c. The lower energy exciton absorption peak at 562 nm (2.21 eV) corresponds to the electronic transitions involving both the ZnSe core and the CdS shell, while the broad exciton peak at 460 nm (2.69 eV) can be attributed to the first excitonic transition in the CdS rod. ZnSe/CdS NRs show intense emission at 592 nm associated with the type-II relaxation of the charge carriers across the ZnSe/CdS junction.

**Figure 2:**
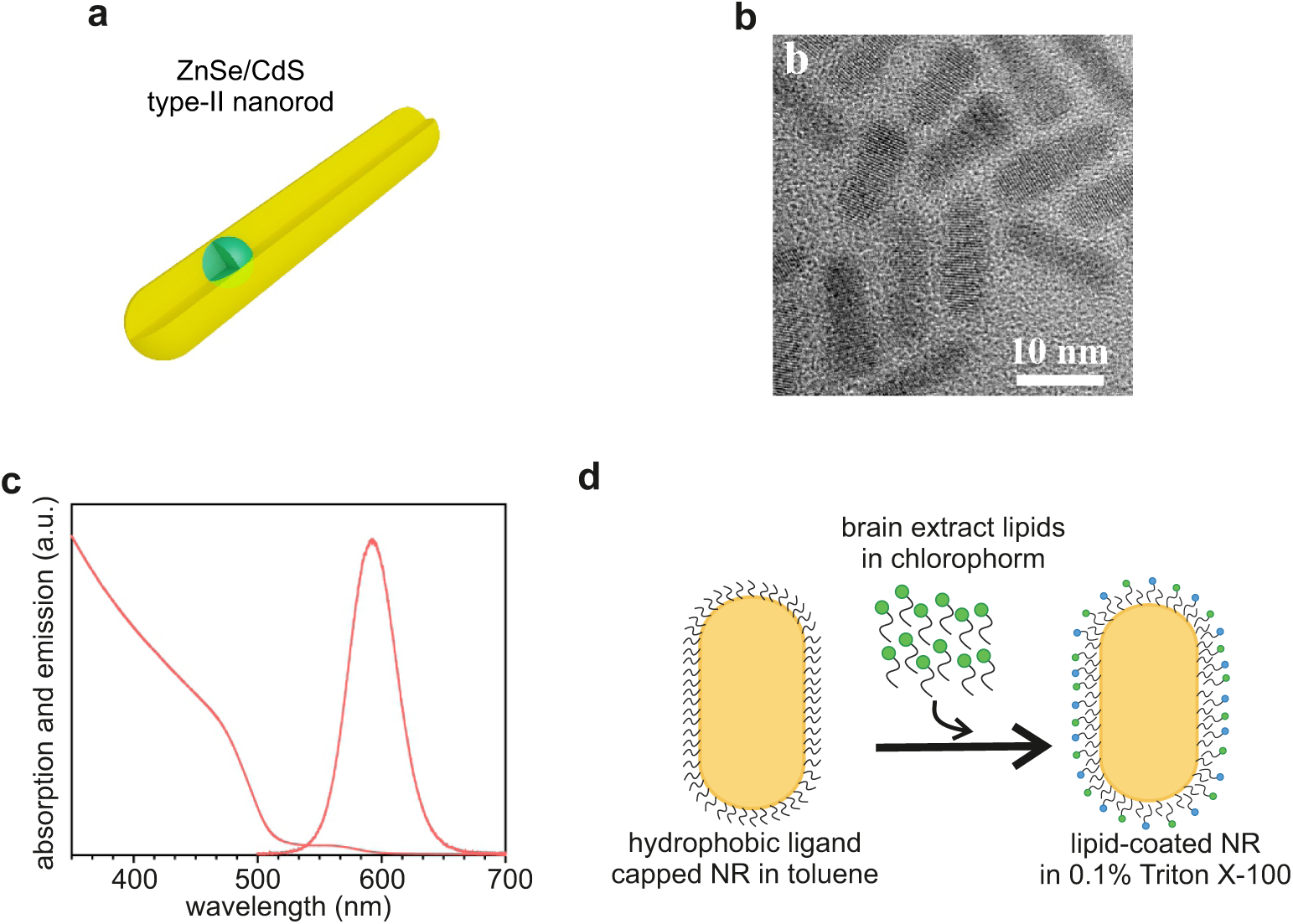
(a) ZnSe/CdS type-II NR schematic. (b) TEM image. (c) Absorption and emission spectra of ZnSe/CdS type-II nanorods. (d) Schematic of NR functionalization.

For voltage sensing in neurons, NRs should be inserted into the neuronal plasma membrane.^37^ After chemical synthesis, the particles unsuitable for the direct use in biological systems. Previously we have shown that surface functionalization with rationally-designed peptides facilitates the insertion of NRs in the plasma membrane of HEK293 cells.^56^ In contrast to HEK293 cells, incorporation of peptide-coated NRs into neurons was insufficient (data not shown), possibly due to differences in membrane composition between the two cell types. As an alternative coating strategy, we tested NRs functionalization with lipids from total brain extract (Fig. 2d, see also Methods, section “Lipid coating”.). Since surface functionalization might change spectral properties of NRs, we measured spectra of 20 individual lipid-coated NRs (see Supporting Information (SI) section “Spectra of single/small cluster vsNRs”). The emission spectrum of the functionalized NRs is centered at 590 nm with a spread of 45 nm (Fig. S1) and it is not different from the spectrum of uncoated NRs (Fig. 2c). We refer to functionalized NRs as vsNRs.

In order to investigate the capacity of vsNRs to label neurons, we diluted the vsNR stock solution in the imaging buffer and deposited directly on neurons cultured on glass coverslips (see Methods section “vsNR and BeRST loading in neuronal culture” for details). When vsNRs were added to the culture, they rapidly attached to the surface of cultured cells (Fig. 3a). We observed dense labeling of neuronal membranes already within a few minutes after the addition of vsNRs to the bath. Fig. 3a left panel shows a bright-field image of the culture, whereas the right panel depicts the matching fluorescent image. Each bright dot on the fluorescent image corresponds to either an individual vsNR or a cluster formed by several vsNRs.

**Figure 3:**
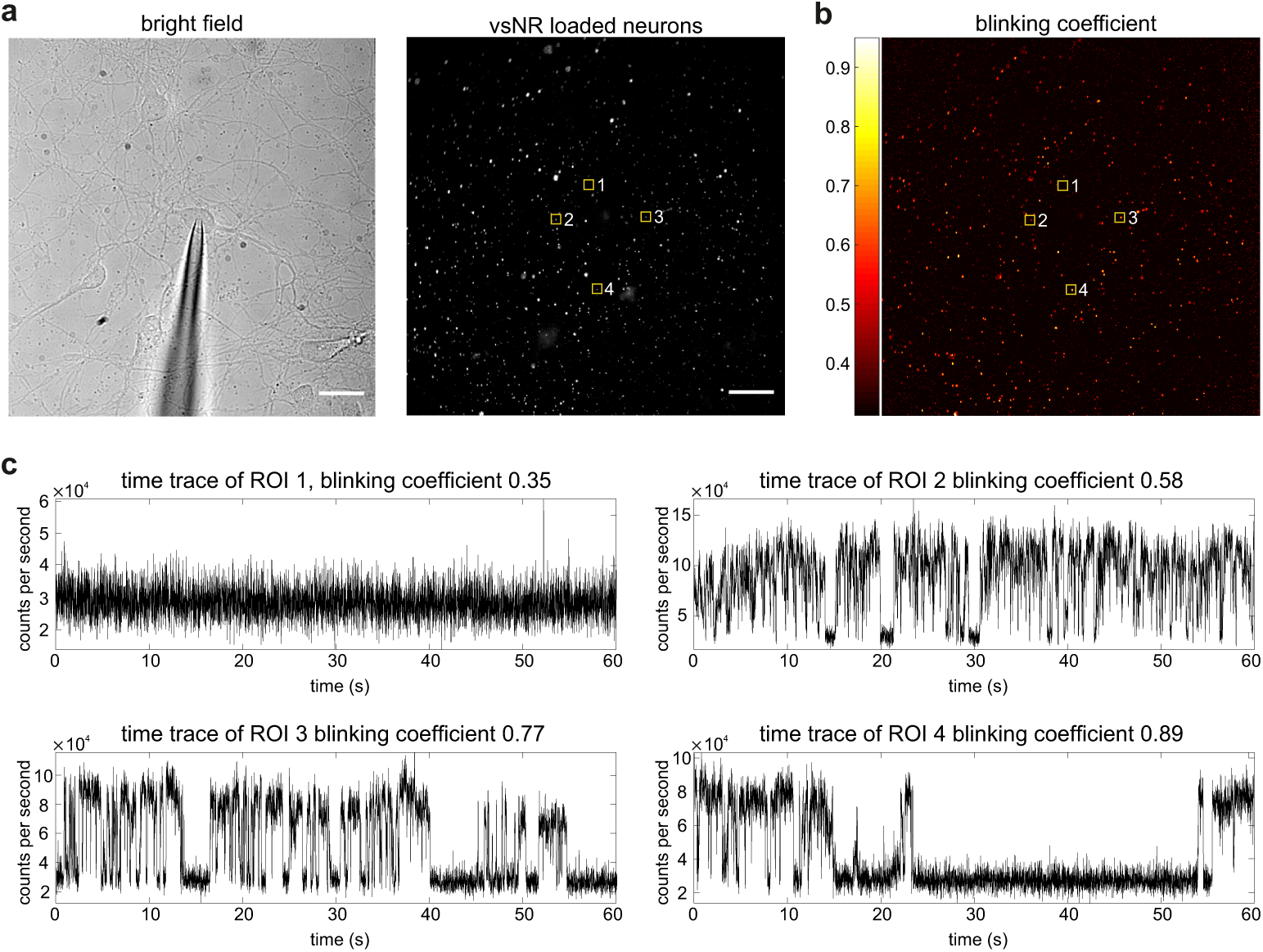
(a) Bright-field (left) and fluorescent (right) images of vsNR-loaded neurons. (b) Pseudo-color image of the blinking coefficient, the same field as in (a). (c) PL time-traces collected from ROIs 1-4, highlighted in (a) and (b) with yellow squares. Scale bars in (a) are 20 *µ*m.

Overall, we found that vsNRs are sufficiently water-soluble to reach the membrane of cultured cells without significant agglomeration, however, they are lipophilic enough to adhere to the membrane and remain stably attached for the time of recording. Importantly, the displacement of adhered particles during recording time did not go beyond the borders of a 1.6×1.6 *µ*m square region used for signal collection. In fact, we found that for the vast majority of vsNR the center of fitted 2d Gaussian point spread function stays inside a single 0.3×0.3 *µ*m pixel at least during one minute recording period (Fig. S2d and SI section “ROI creation and background subtraction”).

### Automated detection of vsNR signal based of blinking properties

We collected intensity from square regions of interest (ROIs) centered at vsNR positions. Custom-made algorithms used for detection of vsNR positions, ROI plotting, and extraction of vsNR signal are described in SI section “ROI creation and background subtraction” and “Blinking analysis and thresholding”. Python and MATLAB custom-made scripts are available in GitHub.^60^

Individual vsNRs can be distinguished from clusters based on the temporal intermittency (blinking) pattern^58^ in their emission (movie S1). The PL intensity distribution of an individual vsNR is multimodal, typically with two states, which we refer to as the “on state” and the “dark state” (Fig. S3a and b), and, possibly, a mixture of them. For bimodal cases, the distance between the mean of each distribution determines the efficiency of the separation between the two states and, as a consequence, the quality of the signal. Blinking of vsNRs in clusters is smeared and does not have pronounced “on” and “dark” states. In order to quantitatively analyze blinking from each time trace, we calculated a blinking coefficient (K_blink_) as follows,

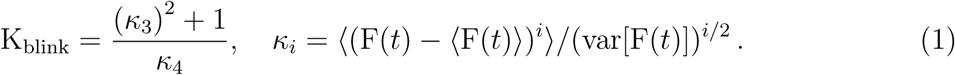

Here F(*t*) is the total PL of the particle. K_blink_ is positively correlated with the distance between the means of the “on state” and the “dark state” intensity distributions of the particle (Fig. S3c). This is due to the properties of the third and the fourth moments of the distribution. The latter goes to small values for broad distributions, such as bimodal ones, and the former increases when they become asymmetric. For example, for a Gaussian distribution, *κ*_3_ = 0 and *κ*_4_ = 3, so K_blink_ = 1/3, while for a distribution with two sharp peaks, *κ*_3_ = 0 and *κ*_4_ = 1, so K_blink_ = 1.

An example of blinking analysis is shown in Fig. 3b and c. Blinking coefficient of each pixel in the field is depicted in pseudocolor (Fig. 3b) and varies from *∼*0.33 for pixels corresponding to background to *∼*0.45 - 0.9 for pixels corresponding to blinking particles. Fig. 3c shows example time traces of 4 ROIs highlighted by yellow squares at Fig. 3a and b. ROI 1 corresponds to a background region, whereas ROIs 2-4 are individual vsNRs with various blinking patterns. Overall, with the blinking coefficient, we can distinguish between individual and clustered vsNRs, and it was used for selection of blinking time traces with a threshold being set at 0.45. A more refined analysis can improve the detection of blinking vsNRs but to the cost of more computational resources or human input. In SI section “Blinking analysis and thresholding”, we study blinking coefficient dependence on the intensity distribution and behavior in randomly selected ROIs.

PL signal collected from a vsNR during its “dark state” corresponds to background noise and, thus, has to be excluded from the recording. To excise “dark states” we calculated a blinking threshold based on the distribution of the overall particle PL intensity (S3b), by fitting it to a sum of two Gaussian distributions, and removed parts of the trace with intensity below the threshold, specifically set by the distribution of the background noise (see SI section “Blinking analysis and thresholding”).

### Imaging and stimulation setup

To test whether vsNRs can sense the membrane potential of neurons we used whole-cell patch-clamp technique. Through the pipette attached to a neuronal body (Fig. 3a, left panel) we modulated the membrane potential of the cell. Fig. 4a depicts the scheme of the setup. For vsNR signal acquisition, we used the left optical port of the microscope equipped with a CMOS camera (left side of Fig. 4a). vsNR PL was excited at 410 nm and collected through 596/83 nm filter to the OptoSplit II dual emission image splitter. Inside the OptoSplit we installed a dichroic beamsplitter with the edge at 590 nm that was used to divide vsNR PL in two parts, roughly at the maximum emission wavelength (Fig. 4b). After the OptoSplit II both parts of the emission signal were projected to the CMOS camera sensor. Camera acquisition at 100 Hz was synchronized with the stimulation (4 frames per period). An example of the split emission image is shown in Fig. 5a, where the left and the right sides of the image represent respectively reflected (*<*590 nm) and transmitted (*>*590 nm) channels collected from the same field. Recording split emission of vsNRs allowed us to assess simultaneously voltage-related fluctuations of vsNR PL intensity (∆F; the sum of the two channels) and voltage-related shift of the vsNR maximum emission wavelength (∆Rt; the ratio between the two channels).

**Figure 4:**
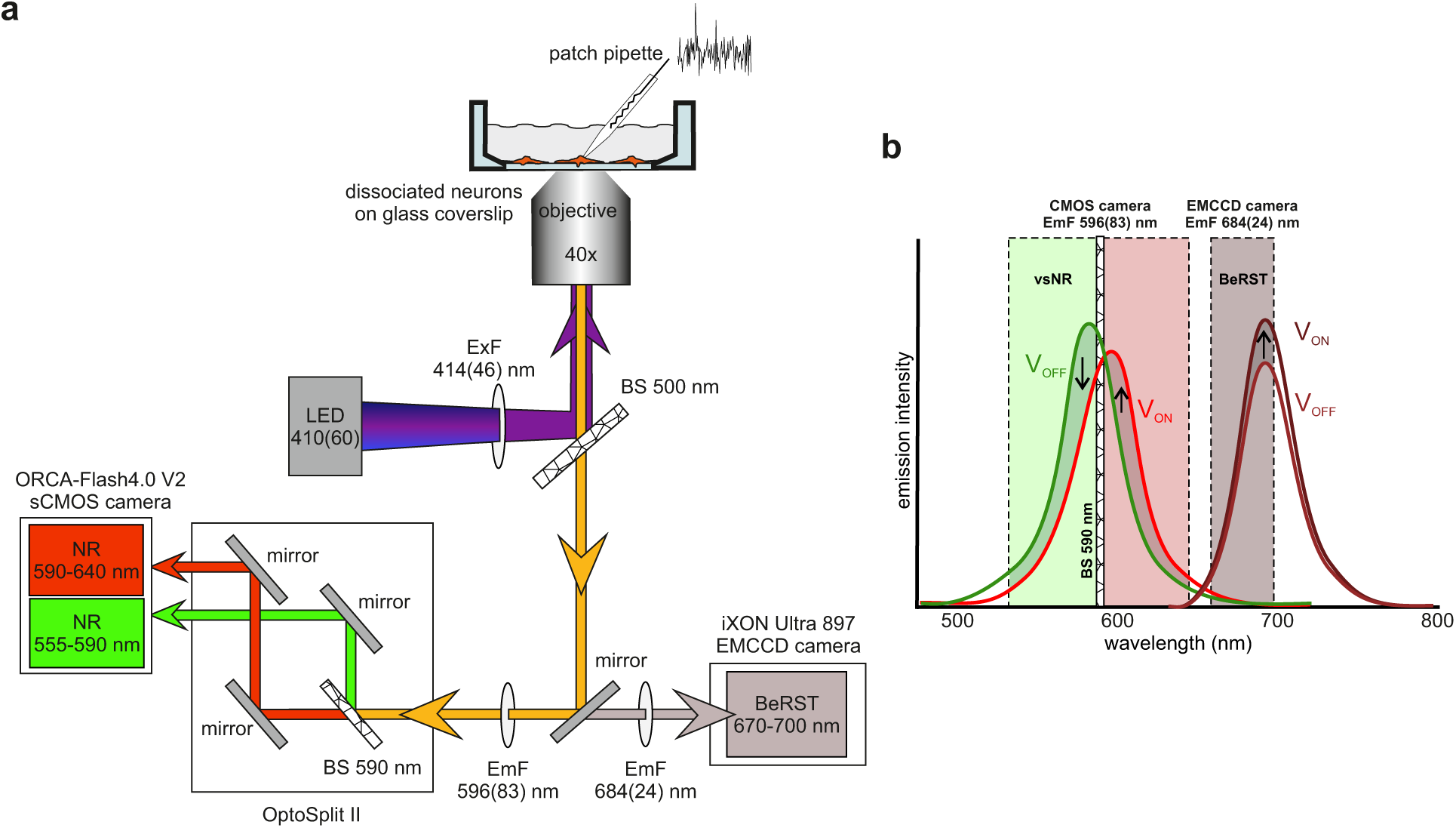
(a) Scheme of the imaging setup. (b) Scheme of the fluorescent channels used for the acquisition of vsNR and BeRST signals.

**Figure 5:**
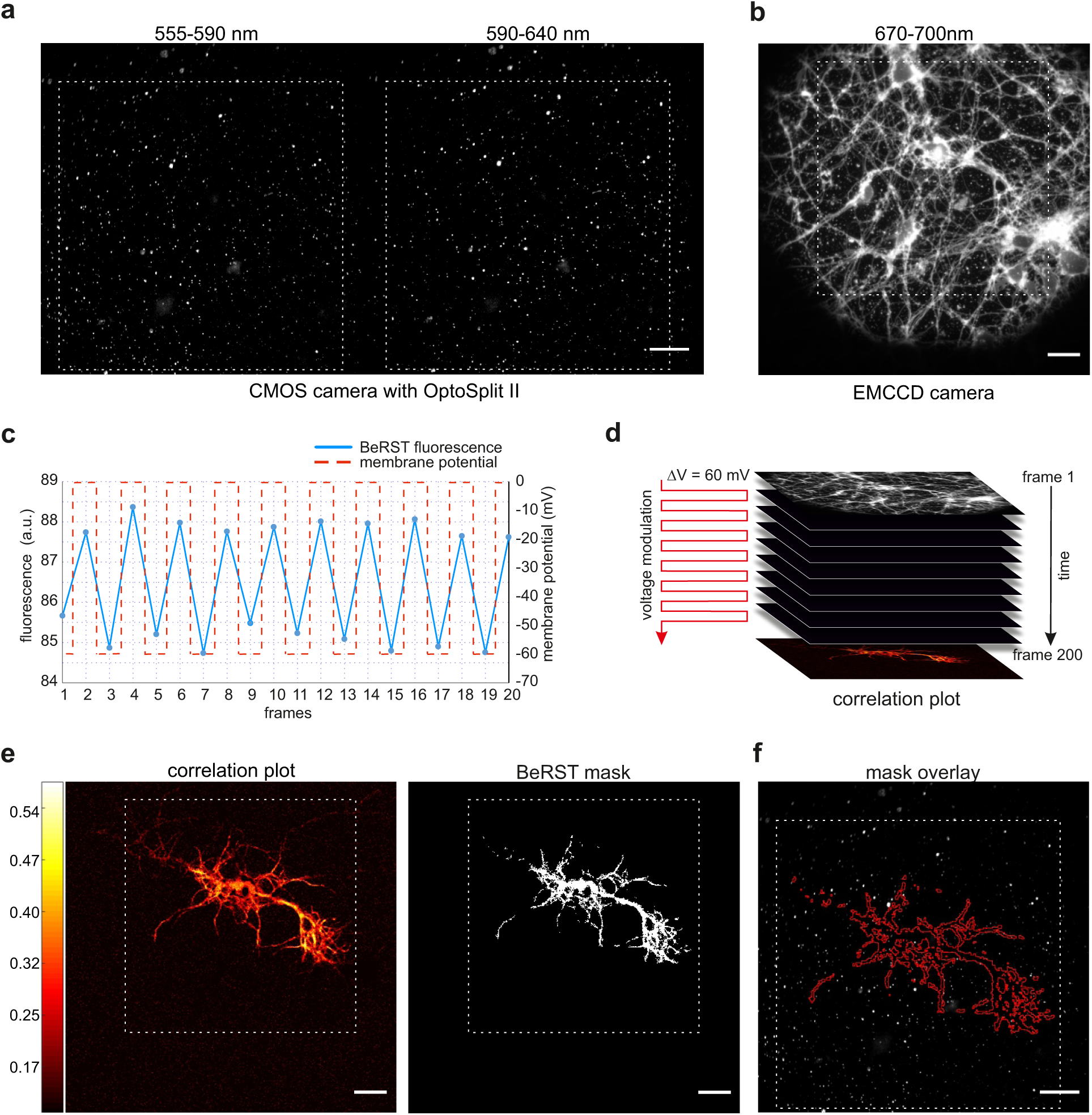
(a) Split emission image of vsNR-loaded neurons. (b) BeRST-loaded neurons, the same field as in (a). (c) BeRST fluorescence time trace (blue line) and membrane voltage modulation (red line). (d) Scheme of generating a plot of responsive pixels, based on the correlation of the BeRST signal with membrane voltage modulation. (e) Pseudo-color plot of the Pearson coefficient for per-pixel temporal correlation between BeRST fluorescence and membrane voltage modulation (left) and the mask produced by thresholding the correlation plot (right). (f) vsNR-loaded neurons with the mask overlay, the same field as in (a) and (b). In (a), (b), (e), and (f) dashed white squares indicate identical fields, scale bars are 20 *µ*m.

### Identification of vsNRs located on stimulated cell

In parallel with vsNRs, we loaded neurons with the voltage-sensitive dye BeRST.^57^ BeRST possess relatively narrow emission spectrum with the maximum at *∼*700 nm that allowed for efficient spectral separation of vsNR and BeRST signals (see SI section “Spectral separation between vsNR and BeRST channels”, Fig. 4b and Fig. S4). BeRST was excited at 400 nm, the emission was filtered through 684/24 nm single-band bandpass filter and collected by iXON Ultra 897 (Andor) EMCCD camera installed at the right optical port of the microscope (right side of Fig. 4a). BeRST displays bright, membrane-localized fluorescence (Fig. 5b) highly sensitive to membrane potential modulation (Fig. 5c).

For each field of view, we performed three optical recordings: two of vsNRs PL intensity, with and without voltage modulation, and one of BeRST fluorescence with voltage modulation. For each pixel of the BeRST channel field, we calculated the Pearson coefficient for the temporal correlation between BeRST fluorescence intensity and voltage modulation applied to the patched neuron (Fig. 5d and Fig. 5e, left panel). After thresholding the coefficient, we obtained a mask that identified the exact pixels in the field of view that were most sensitive to electrophysiological stimulation, typically those that belong to the plasma membrane of the stimulated neuron (Fig. 5e, right panel). Subsequently, we transferred the mask to the vsNR field and used this mask to identify vsNRs located in the correct position, i.e. at the surface of a stimulated cell (Fig. 5f). Of note, the mask corresponds to an underestimated set of responsive pixels, since off-peak excitation of BeRST did not provide enough quantum yield to detect voltage-dependent fluorescence changes in weakly labeled thin processes, due to insufficient signal to noise ratio by single pixels.

### Fourier Transform score

Membrane potential of patched neurons was modulated using a 25 Hz square wave with a 60 mV amplitude. This amplitude roughly corresponds to 150 kV/cm electric field, assuming 4 nm membrane thickness. It is similar or smaller than the stimulation that was used in previous studies (150 mV^56^, 125 - 400 kV/cm^54^, 400 kV/cm^55^), and it is comparable to sub-threshold membrane potential fluctuations that exist in neurons.

Frequency analysis of vsNR signal from stimulated cells revealed a band of 25 Hz in some of the analyzed particles. In order to compare the power of 25 Hz band for different vsNRs in an automated way, we introduced a Fourier Transform (FT) score calculated as the value at 25 Hz divided by the noise of a bandwidth broad enough to account for fluctuations (for more details see SI section “FT score and bootstrap”)

An example of FT score analysis for one field of view is shown in Fig. 6a and b. FT scores for all particles found in the field of view shown in Fig. 6a, left panel, are depicted as a color-coded image in Fig. 6a, right panel. Power spectra of three selected particles marked with yellow squares in Fig. 6a are shown in Fig. 6b. FT score for these particles varies from 3.5, corresponding to 25 Hz band almost indistinguishable from the noise at other frequencies, to 6, corresponding to 25 Hz band that is clearly distinguishable from the noise.

**Figure 6:**
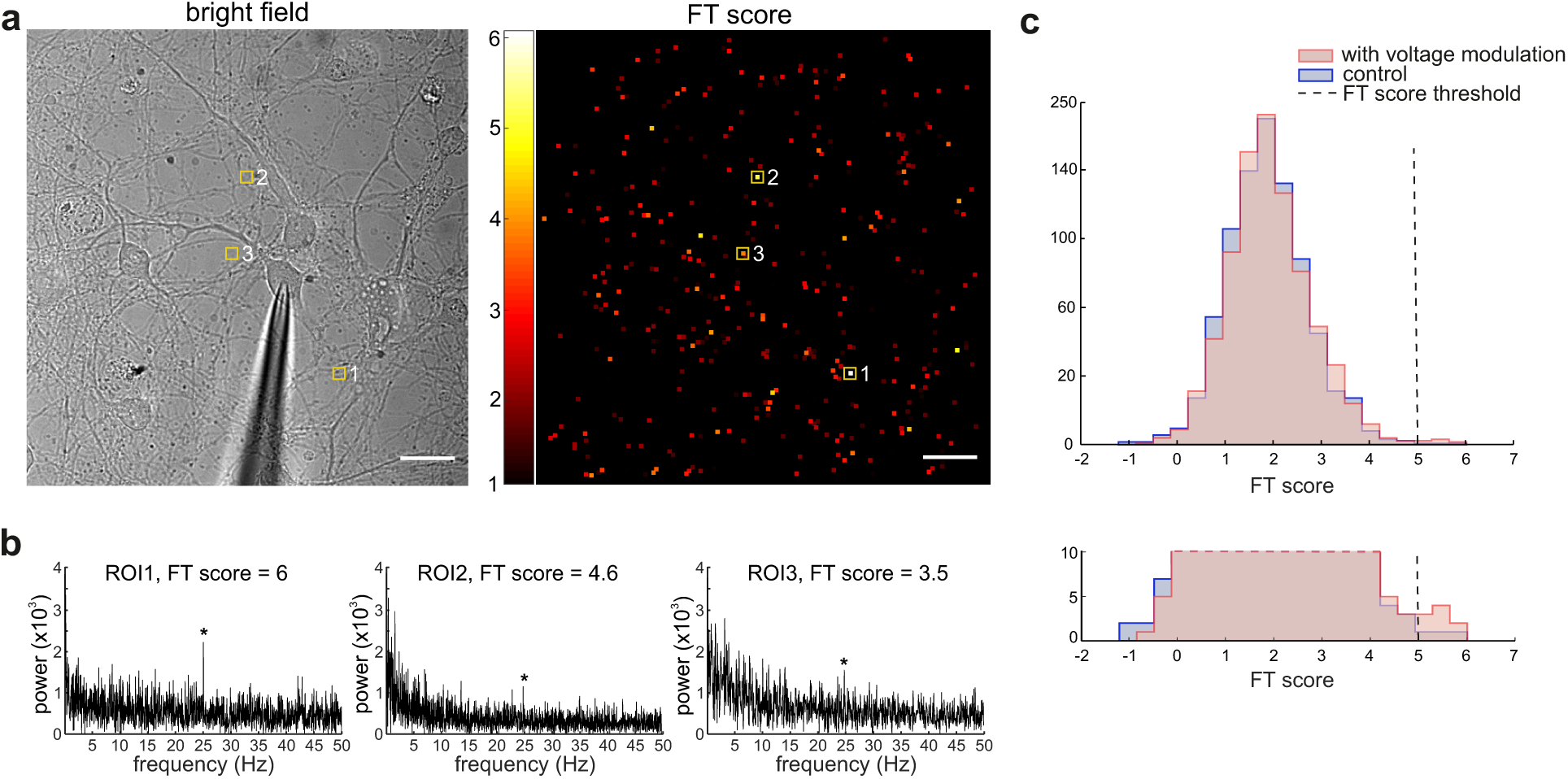
(a) The bright-field image of vsNR-loaded neurons (left panel) and the pseudo-color image of FT score (right panel, the same field as the left panel). (b) Examples of ROIs’ power spectra with different FT score indicated on top of each plot. These ROIs are highlighted in (a) with yellow squares. (c) Histogram of the FT score distribution. 1242 vsNRs were recorded with (pink) and without (blue) voltage modulation. The bottom panel depicts the enlarged fragment of the histogram. Dashed lines indicate the FT score threshold used for vsNR selection.

Altogether we analyzed traces of 1242 particles from 35 cells in 6 independent experiments. The distribution of FT scores of all analyzed particles is shown in Fig. 6c. The mean of FT score for recordings with voltage modulation was not much different from the one for control recordings (with voltage modulation: mean 1.95, standard deviation 0.88; control: mean 1.89, standard deviation 0.85). However, in recordings with voltage modulation, there were significantly more vsNRs with FT score above 5 (Fig. 6c, lower panel). To estimate a probability of getting high FT score by chance, we performed bootstrap analysis of 10^5^ synthetic traces that reproduced the statistical features of vsNR traces in control recordings (see SI section “FT score and bootstrap”). Analysis of control recordings and synthetic traces revealed that variations of vsNR PL unrelated to voltage modulation are high enough to produce a distinguishable band at 25 Hz in power spectra of some traces. Based on the results of the bootstrap analysis we plotted FT score vs the probability of getting this score by chance (Fig. S5c). The probability of getting FT score above 5 is lower than 10^*−*3^ (0.1%). Overall there were 8 vsNRs with FT score of 5 or higher in recordings with voltage modulation (0.6%) and 2 vsNRs in control recordings (0.16% Fig.6c, lower panel). The probability of at least one being a genuine responding vsNRs is 0.9985 (see SI section “FT score and bootstrap”). These particles were selected for further analysis.

We found that only 0.6% of recorded vsNRs contained significant 25 Hz band in the power spectrum of their signal. The rest of the particles were either not responsive, or had very short periods of sensitivity that were below the detection limit of our analysis. Such low success rate seems to be an attribute of live cell recordings since when similar NRs are deposited on a glass and subjected to the external electric field, the proportion of responsive particles is much higher.^54,55^ The major difference between the two conditions is a membrane insertion step. QCSE of an inserted NR is maximized when the particle is oriented perpendicular to the membrane plane with both ends extending out of the membrane on both sides.^46^ Plausibly, in live-cell recordings, non-responsive particles are those that were not properly inserted and are either attached to the membrane surface without transversing it, or misaligned with respect to the membrane plane.

### Burst analysis to detect responsive intervals

For each of the selected responsive particles, we performed the analysis of the trace aimed to reveal voltage-dependent changes in quantum yield and the maximum emission wavelength. For that, we excised blinking periods from PL traces for green (*<*590 nm) and red (*>*590 nm) channels. To preserve the phase of the stimulation square wave, any cycle that contained at least one frame with a signal below the blinking threshold was fully excised. Resulting aligned traces were used to produce F(*t*) and Rt(*t*) as follows:

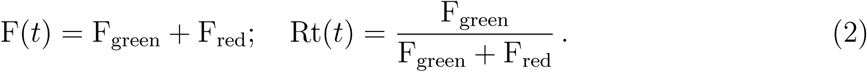

In these calculations F(*t*) is the total PL of the particle and the measure of the particle quantum yield, whereas Rt(*t*) is the proportion of the green channel in total PL that is directly related to the maximum emission wavelength.

The acquisition rate was 4 times the frequency of the stimulation square wave. Thus, each cycle consisted of two frames corresponding to −60 mV and two frames corresponding to 0 mV. We define new variables ∆F*/*F and ∆Rt by averaging frames with identical voltage in each cycle and calculating the difference between −60 mV and 0 mV steps. ∆F was additionally normalized to F_-60mV_ in each cycle to be able to compare arbitrary units of PL intensity between different vsNRs:

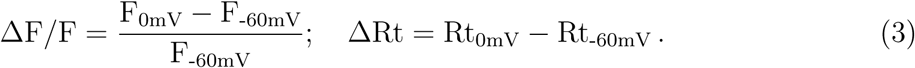

Figure 7a depicts normalized distribution of ∆F*/*F (left panel) and ∆Rt (right panel) values for one recording session of a single particle. These distributions show that during voltage stimulation (red line), but not in the control recording (blue line) there was a number of stimulation cycles with ∆F*/*F *>* 0.5 and also some cycles with ∆Rt *>* 0.1. To determine whether these cycles were distributed randomly across the trace or belonged to some kind of responsive intervals, we followed the approach of a previous work^56^ with some modifications and performed burst-search analysis to identify periods of the trace with pronounced ∆F*/*F or ∆Rt responses. Details of the analysis are described in SI section “Burst intervals” and Fig. S6 that depicts raw and processed time traces of the same particle as in Fig. 7. For this particle, our burst-search algorithm identified 5 periods of pronounced ∆F*/*F (Fig. 7c), all positive, and 7 periods of pronounced ∆Rt (Fig. 7d), 6 positive and one negative (the interval 2). Importantly, ∆F*/*F and ∆Rt intervals significantly overlapped in time (fig.7e). Mean ∆F*/*F for selected intervals was 0.10*±*0.01, whereas mean ∆Rt was 0.012*±*0.006 (fig.7b) This corresponds to roughly 0.6 nm shift of the maximum emission wavelength (SI section “Wavelength shift estimation”). During responsive periods, ∆F*/*F systematically reached 40% for a 60 mV voltage step (for example intervals 1, 2, or 4 in fig.7c). ∆Rt values during responsive intervals could reach 0.1 that corresponds to approximately 5 nm shift of maximum emission wavelength. Another example of single vsNR burst-search analysis is shown in Fig. S7. For that particle six intervals with pronounced ∆F*/*F were identified (mean ∆F*/*F 0.04*±*0.003), whereas no ∆Rt intervals were found.

**Figure 7:**
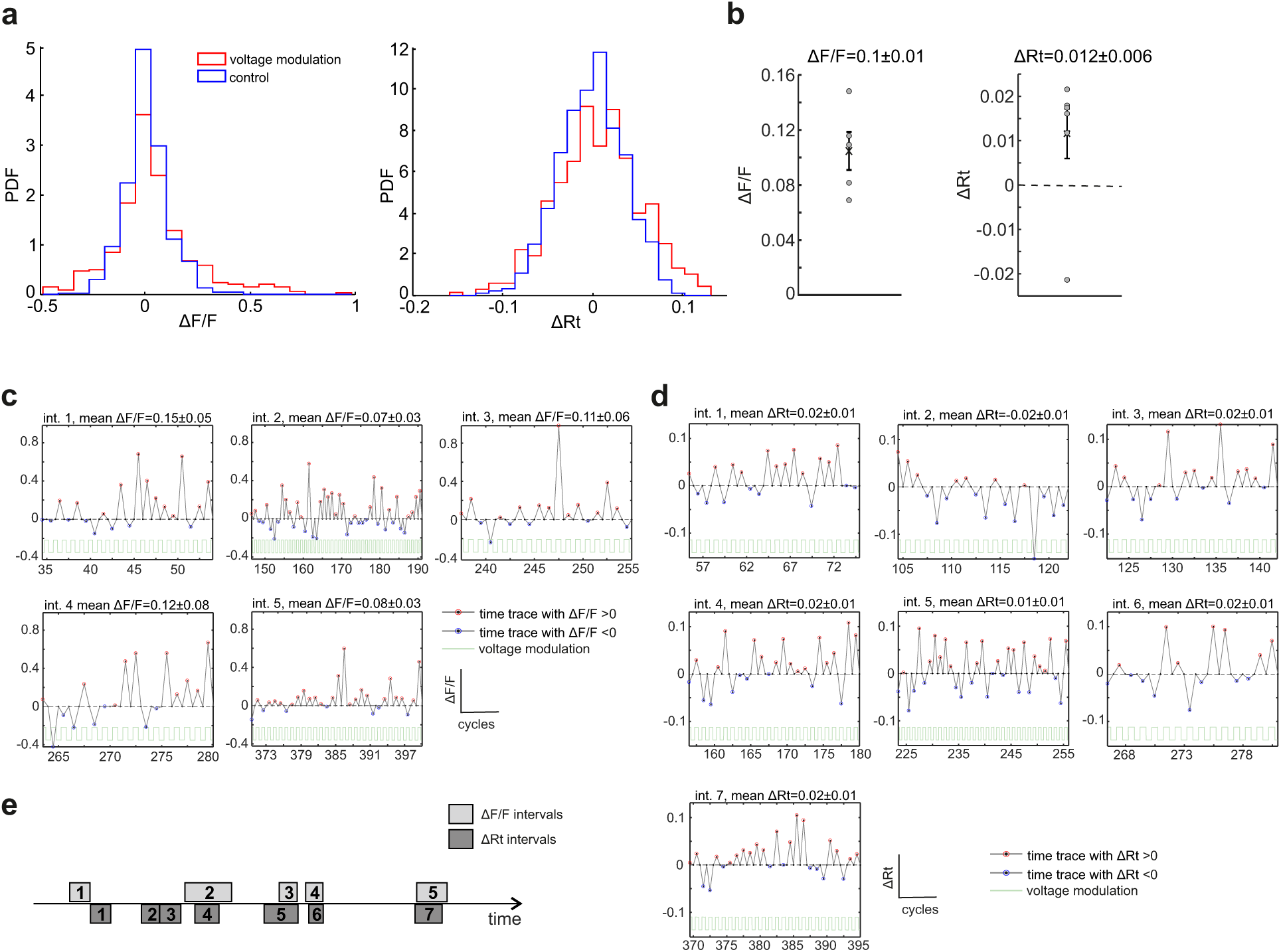
The figure depicts data from a single vsNR located at a neuron and recorded with and without voltage modulation. (a) Distribution of ∆F/F (left panel) and ∆Rt (right panel) of the vsNR with (red) and without (blue) voltage modulation. (b) Mean ∆F/F (left panel) and ∆Rt (right panel) of intervals selected from the trace with voltage modulation. The mean of all selected intervals is indicated with the cross, error bars depict STE. (c) All selected intervals of ∆F/F time trace. (d) All selected intervals of ∆Rt time trace. (e) Relative positions of ∆F/F and ∆Rt intervals over recording time.

Analysis of responsive vsNRs demonstrated that their PL signal exhibited brief periods of voltage sensitivity that were separated by intervals with weak or no voltage response. Previously, we found similar sensitivity bursts in the signal of peptide-coated NRs. ^56^ In the same study, TEM analysis showed that only 16% of peptide-coated NRs were perpendicular to the membrane of small unilamellar vesicles (SUVs), whereas the rest were either tilted or incompletely inserted. These results may reflect not only a steady-state position of a particle but also its temporal fluctuations. In this case, sensitivity bursts could be attributed to the brief periods of proper membrane insertion of the particle. Other reasons for voltage sensitivity fluctuations could be related to the change in the photophysical properties of particles over the time of recording, caused by, for example, ionization of particles,^36^ or temporal fluctuations in particles’ intramembrane neighborhood.

Indeed, despite extensive attempts,^61^ insertion and alignment of particles in the plasma membrane remains problematic. A recent study circumvented this problem by conjugating QDs with fullerene through a peptide linker, ^62^ so that QDs are anchored to the outer leaflet of the plasma membrane, whereas the fullerene is submerged in lipid bilayer. In such configuration the PL of the QDs is quenched in a membrane-potential-dependent manner through electron transfer (ET) from QDs to fullerene, resulting in 7% of depolarization-induced ensemble PL decrease. In contrast, here we show that membrane-integrated QCSE-based sensors, such as vsNRs, during short bursts of voltage sensitivity can exhibit 5-10% of depolarization-induced PL change at a single-particle level. Single-particle recordings potentially allow for super-resolution voltage imaging from confined subcellular locations and gives nanosensors an ultimate advantage over voltage dyes and protein-based sensors. Thus, despite problems with stability in the membrane, membrane-integrated QCSE-based sensors are worth further effort to develop.

## Conclusions

This work is a continuation of our effort to develop voltage nanosensors suitable for monitoring membrane potential in living neurons. We show that lipid-coated type II ZnSe/CdS NRs are capable of reporting the membrane potential of cultured cortical neurons. In the signal collected from individual membrane-bound vsNRs, we detected periods of both voltage-related PL changes and shifts of emission wavelength maximum. Strong fluctuations of voltage sensitivity between particles and within individual traces likely reflect unstable membrane insertion of vsNRs,^56^ which remains the major difficulty of the technique. Overall, we report here extensive measurements of neuronal membrane potential with membrane-integrated vsNRs, as well as the experimental and analysis routine allowing for screening of new surface functionalization methods in order to improve NR insertion in the membrane.

## Methods

### vsNR synthesis

ZnSe/CdS core-shell NRs were synthesized following a previously published procedure with some modifications.^63^

#### Chemicals

Hexadecylamine (HAD, 98%, Sigma-Aldrich), diethylzinc (Et2Zn, 1 M solution in hexane, Sigma-Aldrich), cadmium oxide (99.99%, Sigma-Aldrich), sulfur (99.999%, Sigma-Aldrich), selenium (99.999%, Sigma-Aldrich), oleic acid (OA, 90%, Sigma-Aldrich), trioctylphosphine (TOP, 90%, Sigma-Aldrich), dodecylamine (98%, Fluka), trioctylphosphine oxide (TOPO, technical grade, 99% Sigma-Aldrich), 1-octadecene (ODE, technical grade, 90% Sigma-Aldrich), hexadecylamine (HPA, 99%, Sigma-Aldrich), n-octadecylphosphonic acid (ODPA, 99%, PCI), methanol (anhydrous, 99.8%, Sigma-Aldrich), hexane (anhydrous, 99.9%, Sigma-Aldrich), toluene (99.8%, Sigma-Aldrich). All chemicals were used as received without any further purification.

#### Synthesis of ZnSe NCs

9.4 g of hexadecylamine was degassed under vacuum at 120°C in a reaction flask, under argon flow, the mixture was heated up to 310°C. Then a mixture of 1 mL 1.0 M selenium dissolved in trioctylphosphine, 0.8 mL diethylzinc and 4 mL trioctylphosphine was quickly injected. The reaction was continued at a constant temperature of 270°C for 25 min and then cooled to room temperature.

#### Preparation of cadmium and sulfur stock solutions

0.034 M cadmium oleate was prepared by mixing 0.03 g (0.24 mmol) CdO in 0.6 mL oleic acid and 6.4 mL ODE. The solution was heated to 280°C under argon flow with rigorous stirring until all of the CdO dissolved. 0.29 M S solution was prepared by adding 23.3 mg sulfur in 2.5 mL of dodecylamine at 40°C.

#### CdS shell synthesis

For typical CdS shell coating, 1.1 g unprocessed ZnSe cores, 5.3 mL octadecene (ODE) were loaded into a 50 mL three-neck flask. The solution was degassed at 100°C. After that the solution was heated to 240°C under argon, a mixture of 0.6 mL of 0.034 mmol/mL cadmium oleate stock solution and 0.06 mL of 0.29 mmol/mL sulfur stock solution was injected continuously at 0.72 mL/h. After the injection was finished, the mixture was further annealed for 5 min at 240°C and then cooled down to room temperature. **ZnSe/CdS-CdS NRs Synthesis:** This synthesis was adapted from the previously reported procedure in the literature.^59,64^ In a typical synthesis CdO (60 mg), ODPA (290 mg) and HPA (80 mg) are mixed in TOPO (3.0 g). The mixture is degassed under vacuum at 150°C for 90 min. After the degassing step, the solution was heated to 380°C under argon until it became clear, then 1.8 mL of TOP was injected and the temperature was recovered to 380°C. Subsequently, a solution of 120 mg S in 1.8 mL TOP mix with 40 nmol ZnSe/CdS nanocrystals is rapidly injected. Then the growth was stopped after nanorods grow for 8 min at 365°C. The NRs were precipitated with methanol and dispersed in toluene.

TEM images were taken on a JEOL 2100 TEM equipped with a LaB6 filament at an acceleration voltage of 200 kV.

### Lipid coating

#### Materials

Lipid brain extract (cat# 131101) was purchased from Avanti Polar Lipids. Toluene, Chloroform and Triton X −100 were purchased from Sigma Aldrich.

#### NR surface functionalization by lipid coating

Lipid coating of hydrophobic QDs and NRs is based on weak interactions of intercalating surface ligand alkane chains with the hydrophobic tail of phospholipids. In a typical preparation, a 100 *µ*l solution of 1 *µ*M NR in toluene was mixed with 150 nmol brain extract lipid mixture in chloroform to obtain a total lipid:NR ratio of 1500:1. The organic solvents were evaporated under vacuum at 80°C for 30 min using a benchtop vacuum concentrator (CentriVap, Labconco). The thin film that formed on the vial glass walls was resuspended in 500 *µ*l of 20 mM Tris buffer, 0.1% Triton X-100, pH 7.4. The addition of a lipid detergent was found to significantly improve particle solubility. The sample was then sonicated for 15 min in a bath sonicator (Elmasonic S60H, Elma) at 50°C followed by centrifugation at 20,000 RCF for 2 minutes at 25°C. Particle size distribution and colloidal stability were evaluated by dynamic light scattering measurements from a solution of dispersed particles at 25°C (Zetasizer Nano ZS, Malvern). For a typical preparation, we obtain a single size population with a mean particle hydrodynamic diameter of 14*±*3 nm. This value is used as an indication that the sample consists of a single population of monomeric, non-aggregating, single particles. It is in good agreement with particle dimensions determined by TEM imaging analysis of the particles (Fig. 2b) considering that DLS data analysis yields an average hydrodynamic diameter for non-spherical particle as in the case of vsNRs. Sample solutions stored at 4°C were found to be stable for over a month by routine DLS measurements.

### Dissociated neuronal culture

All animal experiments were approved by the local ethics committee for animal research. Dissociated cortical cultures were prepared from embryonic day 17 mice and cultured for up to one week *in vitro*. We used the protocol described in,^65^ with minor modifications. In brief, embryos were removed from an anesthetized mouse, and cortices were dissected. Cells were dissociated by 0.25% trypsin for 15 min at 37°C and plated on poly-dl-ornithine-coated cover glasses (150,000 cells/cm^2^) in Neurobasal medium containing B27 supplement and 0.5 mM l-glutamine (Gibco/Life Technologies). Before plating, the medium was preincubated on astroglial culture for 24 h. Astroglial cultures were prepared according to a previously published protocol.^65^ In brief, dissociated cortical cells were prepared as described above and plated on uncoated plastic surface. Since neurons require specific coating for attachment, the resulting culture did not contain neurons. This method, together with the use of DMEM medium (Gibco/Life Technologies) supplemented with 10% of fetal bovine serum, 100 U/ml penicillin, and 100 g/ml streptomycin, favors glial survival over neurons.

### vsNR and BeRST loading in neuronal culture

vsNRs were loaded on neuronal cultures right before the imaging session. A cover glass with neurons was removed from the culture dish and placed in ice-cold imaging medium (IM), containing 127 mM NaCl, 3 mM KCl, 2 mM CaCl_2_, 1.3 mM MgCl_2_, 10 mM D-glucose, 10 mM HEPES, pH 7.35. vsNR stock was added to neurons at 1:500 dilution and incubated on ice for 10 min. Then cells were washed with ice-cold IM and the cover glass was transferred to the imaging chamber with room-temperature IM. Voltage-sensitive dye BeRST^57^ was added to the imaging chamber at 50 *µ*M at room temperature and washed out after 5 min.

### Imaging and patch clamp

For imaging, we used Nikon eclipse Ti epifluorescence microscope (Fig. 4a) equipped with ORCA-Flash 4.0 V2 (Hamamatsu Photonics) CMOS camera and iXON Ultra 897 (Andor) EMCCD camera, as well as patch-clamp hardware (Axon). Fluorescence was excited by 410 nm LED (Solis, Thorlabs) at 7.5W/cm^2^ for vsNRs or at 0.8W/cm^2^ for BeRST recordings. Emission was collected through Nikon Plan Fluor 40x/1.30 oil objective and passed through 596/83 nm single-band bandpass filter (Semrock FF01-596/83-25) to the OptoSplit II dual emission image splitter for vsNRs, or through 684/24 nm single-band bandpass filter (Semrock FF02-684/24-25) for BeRST. Inside the OptoSplit we installed a dichroic beamsplitter with the edge at 590 nm (Chroma T590lpxr-UF3).

Somatic recordings from cultured neurons were performed at room temperature in the whole-cell voltage-clamp mode using a Multiclamp 700B controlled by pClamp 10 acquisition software (Molecular Devices). Currents were filtered at 2-3 kHz and sampled at 10 kHz using a Digidata 1440 A (Molecular Devices). Patch pipettes (4-6 MΩ) were filled with internal solution containing 130 mM K gluconate, 10 mM KCl, 4 mM Mg-ATP, 0.5 mM Na-GTP, 10 mM phosphocreatine, and 10 mM HEPES, pH 7.4.

For each field, we performed three recordings, two for vsNRs, with and without voltage modulation, and one for BeRST with voltage modulation. For vsNRs recordings, the membrane potential of patched neurons was either modulated using a 25 Hz square wave with 60 mV amplitude or hold at −60 mV (control recording). To synchronize imaging and voltage modulation, each frame of the camera was triggered by Digidata 1440 A at 100Hz, 4 times the frequency of the stimulation square wave. Each recording lasted for 60 sec and contained 6000 frames of time-lapse movie. For BeRST recordings, membrane potential was modulated at 5Hz with 60 mV amplitude, and the camera was triggered at 10Hz.

After the acquisition, time-lapse movies were directly saved in tif format and processed for analysis according to the workflow scheme (Fig. 1) without any further modifications.

### Data analysis

Data analysis was performed both in python and matlab with in-house scripts. Python scripts were arranged in jupyter/ipython notebooks that require some basic scientific libraries (numpy, matplotlib, scipy and pillow) and can be re-used easily. All the software used to produce the figures of this manuscript is released as open-source and is available in GitHub.^60^

## Acknowledgement

The authors thank Xavier Marques and Astou Tangara for the tangible support in setup optimization and maintenance.

This research was supported by the European Research Council (ERC) advanced grant NVS 669941, by the Human Frontier Science Program (HFSP) research grant RGP0061/2015, by the BER program of the Department of Energy Office of Science grant DE-FC03-02ER63421, by the STROBE National Science Foundation Science & Technology Center, Grant No. DMR-1548924, and by the European Unions Horizon 2020 Framework Programme for Research and Innovation under the Specific Grant Agreement No. 785907 (Human Brain Project SGA2). A.L. acknowledges support from the Marie Curie Individual Fellowship NanoVoltSens 752019. G.O. acknowledges HHMI Gilliam Fellows program. E.M. acknowledges NIH support (R35GM119855).

## References

(1) Buzsáki, G.; Anastassiou, C.; Koch, C. Nature Reviews Neuroscience 2012, 13, 407–20.

(2) Lin, M. Z.; Schnitzer, M. J. Nature neuroscience 2016, 19, 1142–53.

(3) Grienberger, C.; Konnerth, A. Neuron 2012, 73, 862–85.

(4) Davila, H. V.; Salzberg, B. M.; Cohen, L. B.; Waggoner, A. S. A large change in axon fluorescence that provides a promising method for measuring membrane potential. 1973.

(5) Loew, L. M.; Bonneville, G. W.; Surow, J. Biochemistry 1978, 17, 4065–4071.

(6) Fluhler, E.; Burnham, V. G.; Loew, L. M. Biochemistry 1985, 24, 5749–5755.

(7) Shoham, D.; Glaser, D. E.; Arieli, A.; Kenet, T.; Wijnbergen, C.; Toledo, Y.; Hildesheim, R.; Grinvald, A. Neuron 1999, 24, 791–802.

(8) Yan, P.; Acker, C. D.; Zhou, W. L.; Lee, P.; Bollensdorff, C.; Negreane, A.; Lotti, J.; Sacconi, L.; Antic, S. D.; Kohl, P.; Mansvelder, H. D.; Pavone, F. S.; Loew, L. M. Proceedings of the National Academy of Sciences of the United States of America 2012, 109, 20443–20448.

(9) Braubach, O.; Cohen, L. B.; Choi, Y. Advances in experimental medicine and biology 2015, 859, 3–26.

(10) Grinvald, A.; Petersen, C. C. Imaging the Dynamics of Neocortical Population Activity in Behaving and Freely Moving Mammals. 2015.

(11) Miller, E. W. Current Opinion in Chemical Biology 2016, 33, 74–80.

(12) Jin, L.; Han, Z.; Platisa, J.; Wooltorton, J. R. A.; Cohen, L. B.; Pieribone, V. A. Neuron 2012, 75, 779–785.

(13) St-Pierre, F.; Marshall, J. D.; Yang, Y.; Gong, Y.; Schnitzer, M. J.; Lin, M. Z. Nature neuroscience 2014, 17, 884–9.

(14) Zou, P.; Zhao, Y.; Douglass, A. D.; Hochbaum, D. R.; Brinks, D.; Werley, C. a.; Harrison, D. J.; Campbell, R. E.; Cohen, A. E. Nature communications 2014, 5, 4625.

(15) Hochbaum, D. R. et al. Nature methods 2014, 11, 825–33.

(16) Gong, Y.; Huang, C.; Li, J. Z.; Grewe, B. F.; Zhang, Y.; Eismann, S.; Schnitzer, M. J. Science 2015, 350, 1361–1366.

(17) Piao, H. H.; Rajakumar, D.; Kang, B. E.; Kim, E. H.; Baker, B. J. Journal of Neuroscience 2015, 35, 372–385.

(18) Abdelfattah, A. S. et al. Science 2019, 365, 699–704.

(19) Knöpfel, T.; Gallero-Salas, Y.; Song, C. Current Opinion in Chemical Biology 2015, 27, 75–83.

(20) Nakajima, R.; Jung, A.; Yoon, B. J.; Baker, B. J. Frontiers in Synaptic Neuroscience 2016, 8, 1–9.

(21) Platisa, J.; Pieribone, V. A. Current Opinion in Neurobiology 2018, 50, 146–153.

(22) Wang, D.; Zhang, Z.; Chanda, B.; Jackson, M. B. Biophysical Journal 2010, 99, 2355–2365.

(23) Liu, P.; Grenier, V.; Hong, W.; Muller, V. R.; Miller, E. W. Journal of the American Chemical Society 2017, 139, 17334–17340.

(24) Grenier, V.; Daws, B. R.; Liu, P.; Miller, E. W. Journal of the American Chemical Society 2019, 141, 1349–1358.

(25) Cao, G.; Platisa, J.; Pieribone, V. A.; Raccuglia, D.; Kunst, M.; Nitabach, M. N. Cell 2013, 154, 904–913.

(26) Flytzanis, N. C.; Bedbrook, C. N.; Chiu, H.; Engqvist, M. K.; Xiao, C.; Chan, K. Y.; Sternberg, P. W.; Arnold, F. H.; Gradinaru, V. Nature Communications 2014, 5.

(27) Kibat, C.; Krishnan, S.; Ramaswamy, M.; Baker, B. J.; Jesuthasan, S. Journal of Neurogenetics 2016, 30, 80–88.

(28) Yang, H. H.; Sun, X.; Ding, X.; Lin, M. Z.; Clandinin, T. R. Cell 2016, 166, 1–13.

(29) Raccuglia, D.; McCurdy, L. Y.; Demir, M.; Gorur-Shandilya, S.; Kunst, M.; Emonet, T.; Nitabach, M. N. eNeuro 2016, 3.

(30) Chen, D.; Sitaraman, D.; Chen, N.; Jin, X.; Han, C.; Chen, J.; Sun, M.; Baker, B. S.; Nitabach, M. N.; Pan, Y. Nature Communications 2017, 8, 1–13.

(31) Chamberland, S.; Yang, H. H.; Pan, M. M.; Evans, S. W.; Guan, S.; Chavarha, M.; Yang, Y.; Salesse, C.; Wu, H.; Wu, J. C.; Clandinin, T. R.; Toth, K.; Lin, M. Z.; St-Pierre, F. eLife 2017, 6, 1–35.

(32) Peterka, D. S.; Takahashi, H.; Yuste, R. Neuron 2011, 69, 9–21.

(33) Kulkarni, R. U.; Miller, E. W. Biochemistry 2017,

(34) Bando, Y.; Grimm, C.; Cornejo, V. H.; Yuste, R. BMC Biology 2019, 17, 1–12.

(35) Marshall, J. D.; Schnitzer, M. J. ACS Nano 2013, 7, 4601–4609.

(36) Rowland, C. E.; Susumu, K.; Stewart, M. H.; Oh, E.; Mäkinen, A. J.; O’Shaughnessy, T. J.; Kushto, G.; Wolak, M. A.; Erickson, J. S.; L. Efros, A.; Huston, A. L.; Delehanty, J. B. Nano Letters 2015, 15, 6848–6854.

(37) Park, K.; Weiss, S. Biophysical Journal 2017, 112, 703–713.

(38) Efros, A. L.; Delehanty, J. B.; Huston, A. L.; Medintz, I. L.; Barbic, M.; Harris, T. D. Nature Nanotechnology 2018, 13, 278–288.

(39) Miller, D. A. B.; Chemla, D. S.; Damen, T. C.; Gossard, A. C.; Wiegmann, W.; Wood, T. H.; Burrus, C. A. Physical Review Letters 1984, 53, 2173–2176.

(40) Empedocles, S.; Bawendi, M. Science 1997, 278, 2114–2116.

(41) Takeuchi, T.; Sota, S.; Sakai, H.; Amanoa, H.; Akasaki, I.; Kaneko, Y.; Nakagawa, S.; Yamaoka, Y.; Yamada, N. Journal of Crystal Growth 1998. 189-190, 616–620.

(42) Scott, R.; Achtstein, A. W.; Prudnikau, A. V.; Antanovich, A.; Siebbeles, L. D.; Artemyev, M.; Woggon, U. Nano Letters 2016, 16, 6576–6583.

(43) Reiss, P.; Protière, M.; Li, L. Small 2009, 5, 154–168.

(44) Becker, K.; Lupton, J. M.; Müller, J.; Rogach, A. L.; Talapin, D. V.; Weller, H.; Feldmann, J. Nature Materials 2006, 5, 777–781.

(45) Kraus, R. M.; Lagoudakis, P. G.; Rogach, A. L.; Talapin, D. V.; Weller, H.; Lupton, J. M.; Feldmann, J. Physical Review Letters 2007, 98, 017401.

(46) Park, K.; Deutsch, Z.; Li, J. J.; Oron, D.; Weiss, S. ACS Nano 2012, 6, 10013–10023.

(47) Nuriya, M.; Jiang, J.; Nemet, B.; Eisenthal, K. B.; Yuste, R. Biophotonics International 2006, 103, 786–790.

(48) Stuart, G. J.; Palmer, L. M. Pflugers Archiv European Journal of Physiology 2006, 453, 403–410.

(49) Palmer, L. M.; Stuart, G. J. The Journal of neuroscience: the official journal of the Society for Neuroscience 2009, 29, 6897–6903.

(50) Acker, C. D.; Yan, P.; Loew, L. M. Biophysical Journal 2011, 101, L11–L13.

(51) Ford, K. J.; Davis, G. W. Journal of Neuroscience 2014, 34, 14517–14525.

(52) Popovic, M. A.; Carnevale, N.; Rozsa, B.; Zecevic, D. Nature Communications 2015, 6, 8436.

(53) Kwon, T.; Sakamoto, M.; Peterka, D. S.; Yuste, R. Cell Reports 2017, 20, 1100–1110.

(54) Bar-Elli, O.; Steinitz, D.; Yang, G.; Tenne, R.; Ludwig, A.; Kuo, Y.; Triller, A.; Weiss, S.; Oron, D. ACS Photonics 2018, 5, 2860–2867.

(55) Kuo, Y.; Li, J.; Michalet, X.; Chizhik, A.; Meir, N.; Bar-Elli, O.; Chan, E.; Oron, D.; Enderlein, J.; Weiss, S. ACS Photonics 2018, 5, 4788–4800.

(56) Park, K. et al. Science Advances 2018, 4.

(57) Huang, Y. L.; Walker, A. S.; Miller, E. W. Journal of the American Chemical Society 2015, 137, 10767–10776.

(58) Nirmal, M.; Dabbousi, B. O.; Bawendi, M. G.; Macklin, J. J.; Trautman, J. K.; Harris, T. D.; Brus, L. E. Nature 1996, 383, 802–804.

(59) Dorfs, D.; Salant, A.; Popov, I.; Banin, U. Small 2008, 4, 1319–1323.

(60) Code library - github repository. 2019; https://github.com/pabloserna/Nanorods.

(61) Molokanova, E.; Bartel, J. A.; Zhao, W.; Naasani, I.; Ignatius, M. J.; Treadway, J. A.; Savtchenko, A. Biophotonics Int. 2008, 2631.

(62) Nag, O. K. et al. ACS Nano 2017, 11, 5598–5613.

(63) Tyrakowski, C. M.; Shamirian, A.; Rowland, C. E.; Shen, H.; Das, A.; Schaller, R. D.; Snee, P. T. Chemistry of Materials 2015, 27, 7276–7281.

(64) Carbone, L. et al. Nano Letters 2007, 7, 2942–2950.

(65) Banker, G.; Goslin, K. Culturing Nerve Cells, second edition; A Bradford Book, 1998.

(66) Chroma T590lpxr-UF3 dichroic beamsplitter. 2019; https://www.chroma.com/products/parts/t590lpxr#tabs-0-main-1.

(67) Thompson, R. E.; Larson, D. R.; Webb, W. W. Biophysical journal 2002, 82, 2775–2783.

(68) Shimizu, K. T.; Neuhauser, R. G.; Leatherdale, C. A.; Empedocles, S. A.; Woo, W. K.; Bawendi, M. G. Physical Review B 2001, 63, 205316.

