## Supplemental Information for "Feasibility analysis of semiconductor voltage nanosensors for neuronal membrane potential sensing"

### Supporting Information Available

Supplementary material for this article includes the following sections: (1) Spectra of single/small cluster vsNRs; (2) Wavelength shift estimation; (3) ROI creation and background subtraction; (4) Blinking analysis and thresholding; (5) Spectral separation between vsNR and BeRST channels; (6) FT score and bootstrap; (7) Burst intervals; (8) movie S1, vsNR-loaded cultured cortical neurons; (9) code library in github repository.[60]

This material is available free of charge via the Internet at `http://pubs.acs.org/`.

### Spectra of single/small cluster vsNRs

Samples were prepared by drop-casting a solution of vsNR on a glass coverslip. They were placed in an inverted microscope and imaged using a high NA (1.4) objective. Individual particles and clusters were illuminated by a 470 nm laser, the PL was collected into a fiber and directed to a spectrometer (SP2300, Princeton instruments). The measured spectra are shown in Fig. S1a. After normalization (Fig. S1b), we extracted the ensemble statistics, both the averaged spectrum (Fig. S1c) and full width at half maximum (FWHM, Fig. S1d). Acquisition time ranged from 5 to 60 sec.

### Wavelength shift estimation

We followed a recent study[54] to estimate the wavelength shift corresponding to the difference in ratio ΔRt. First, we modeled the emission spectrum as a Gaussian distribution with $\mu = 590$ nm and $\sigma = 19.2$ nm. By using the properties of the dichroic beamsplitter installed in the Optosplit II (Chroma T590lpxr-UF3[66]), we calculated the amount of light that goes through each channel and the ratio of both. This value is roughly 0.5 since we used a specific filter to split both channels equally. Then, we shifted the mean parameter of the Gaussian distribution by $\delta$ nm and estimated the ratio. The difference between the two ratios was

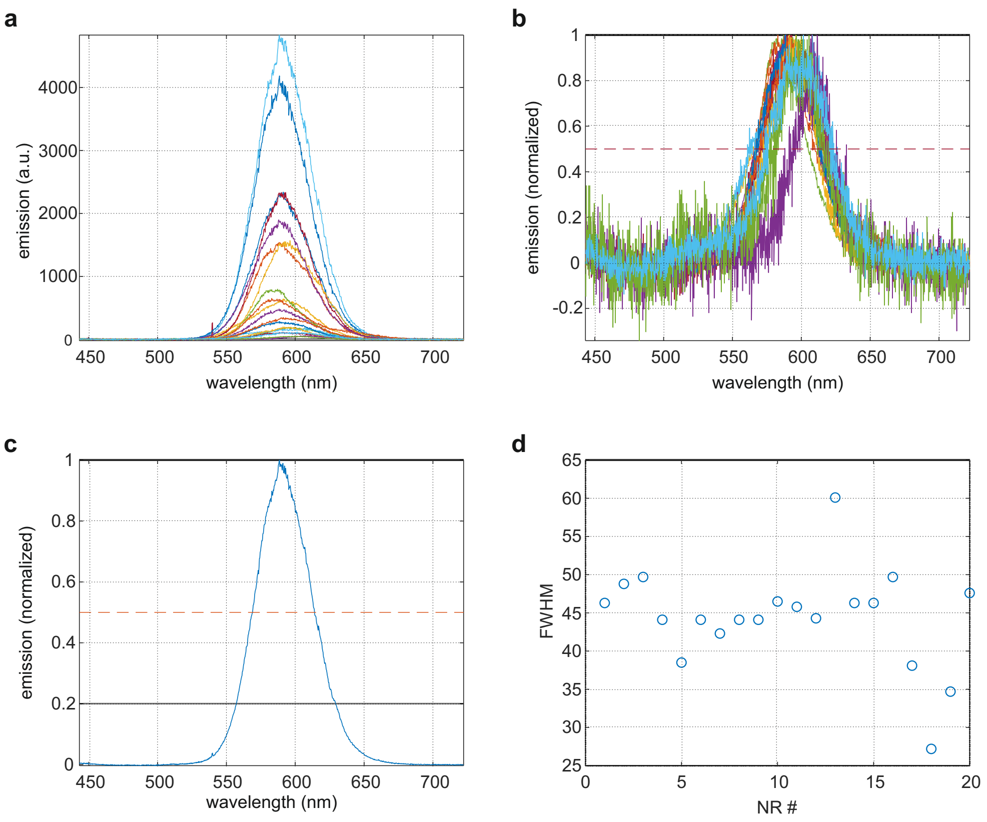


Figure S1: (a) Spectra of 20 single/small cluster vsNRs (b) Normalized spectra of the same vsNRs, as in (a). (c) Mean spectrum of 20 vsNRs (d) Full width at half maximum (FWHM) of the recorded vsNRs.

approximately proportional to $\delta$ for the range $\delta \in (-10\text{nm}, 10\text{nm})$.We used this to estimate the wavelength shift from experimentally measured $\Delta$Rt values.

### ROI creation and background subtraction

**ROI creation.** To obtain regions of interest (ROIs) that include vsNR signal, we first searched for local intensity maxima in the projected (averaged) image. We used a standard algorithm, consisting in a *dilation* of the original picture, i.e., a 5x5 max-pooling with stride 1 and ”same” padding, and compared it with the original image pixel-wise. Those pixels where both images coincide correspond to local maxima and thus putative ROIs.

**Local background subtraction.** We used these putative ROIs to generate time traces by determining for each ROI background intensity and integrating the signal. For a given 5x5 ROI (Fig. S2a) and at each time frame, we sorted the intensity values of the 25 pixels. We set the average fluorescence of the five darkest pixels times 20 as the background of that particular time frame and ROI. The sum intensity of the 20 remaining pixels are referred to as "signal". In Fig. S2b, we show three non-blinking frames of the same ROI, where the background pixels have been marked with white dots, and three frames where the vsNR was in the dark state. In Fig. S2c (upper panel) the signal and background time traces are represented as function of time for this ROI. The final fluorescence time trace is defined by subtracting the background from the signal (Fig. S2c lower panel).

**vsNR membrane mobility.** vsNRs that stay attached to the cell surface after washing coverslips were relatively immobile. We have quantified the mobility of the recorded particles with standard tools in single-particle tracking, using in-house written python scripts. To do this we have fitted each ROI to a general 2d Gaussian Point Spread Function (PSF), except for a rotation which we have checked it was not relevant,

$$P(x, y) = \frac{1}{2\pi\sigma_x\sigma_y} \exp\left(-\frac{(x-\mu_x)^2}{2\sigma_x^2} - \frac{(y-\mu_y)^2}{2\sigma_y^2}\right) \quad , \tag{4}$$

where $\sigma_x$ and $\sigma_y$ are the standard deviations in directions $X$ and $Y$ respectively, and $\mu_x$ and $\mu_y$ the center of the ROI.

The distribution of $\mu_x$ and $\mu_y$ for the example ROI is shown in Fig. S2d. Remarkably, the fitted position of the center of the ROI stayed within a single 0.3x0.3 $\mu$m pixel during the entire movie (1 min). This was the case for the vast majority of ROIs we have studied here. It is also interesting to compare this position with the localization precision due to both pixel size and background noise. Following,[67] the localization precision is affected by two terms, one dependent on the pixel size $a$ and the other dependent on the signal to noise

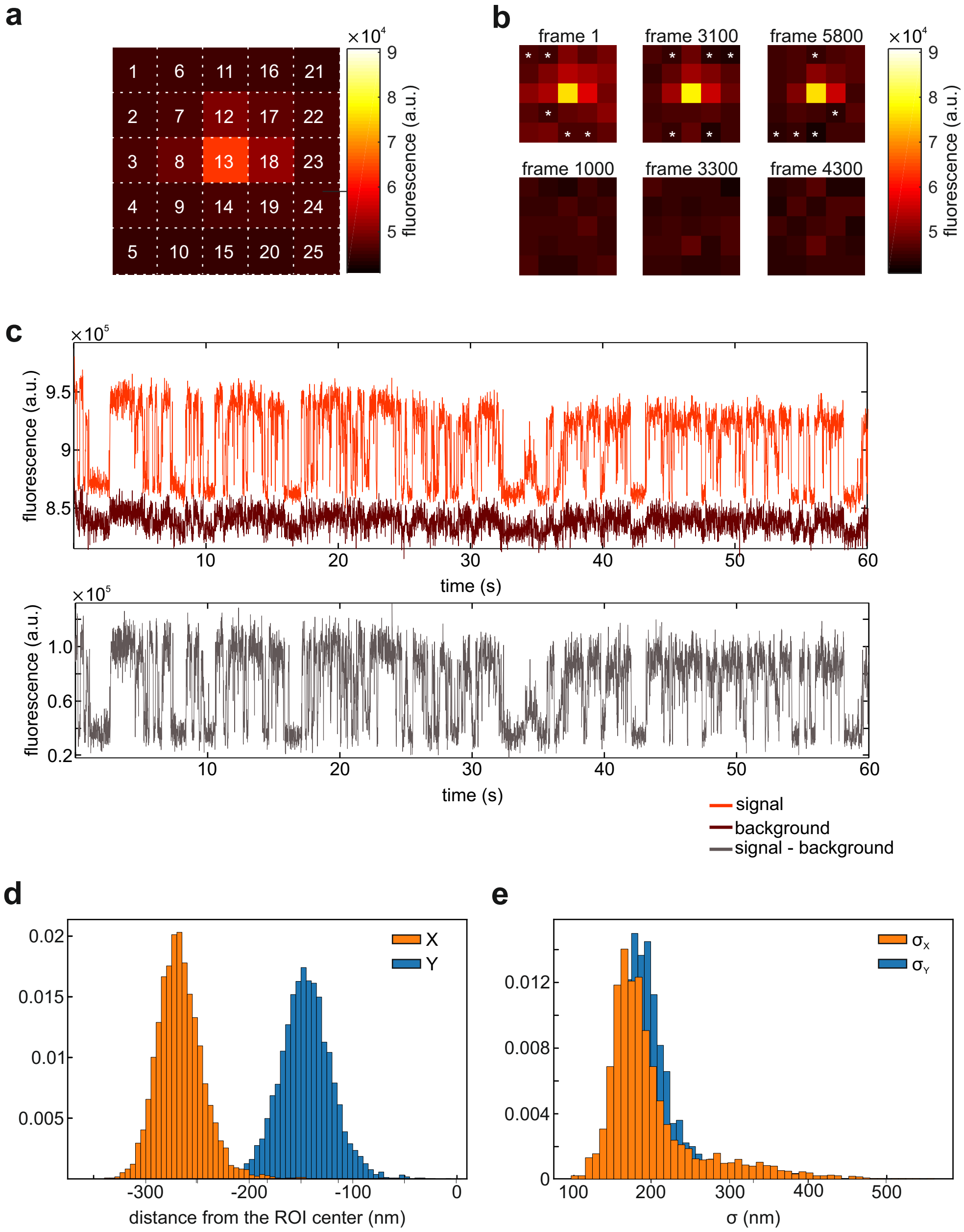


Figure S2: (a) An example of 5x5 pixels ROI with a vsNR in the center. Color-coding refers to the mean intensity of each pixel over the entire recording time (60 s). (b) Selected time points from the recording as in (a). Upper row images depict frames during "on" state of the vsNR, whereas bottom row images show frames from blinking periods. White asterisks in the upper row images indicate pixels with minimal intensity selected for background calculation. (c) Fluorescent time trace of the same vsNR as in (a) and (b). The upper panel depicts the signal (light red) and the background (dark red), whereas the bottom panel shows the signal with the subtracted background. (d) Distribution of the center position of the fitted PSF, defined in Eq. 5, from the ROI center for a period of 1 minute, without considering blinking periods. (e) Distribution of the width of the PSF for the ensemble of recorded vsNRs.

ratio,

$$\langle(\Delta x)^2\rangle = \frac{\sigma_x^2 + a^2/12}{N} + \frac{8\pi\sigma_x^4 b^2}{a^2 N^2} \quad , \tag{5}$$

where $b$ is the background noise per pixel and $N$ the number of photons measured. For the particular example shown in Fig. S2, $\sigma_x \approx 191$ nm, and using the parameters of our setup, the localization precision was roughly 15 nm. This value is smaller than the standard deviation of the distribution of positions shown in panel d. In panel e, we show the distribution of $\sigma_x$ and $\sigma_y$ for the particles we have studied. The distribution was skewed, partly due to heterogeneity of the environment and response of vsNRs, but we cannot discard the presence of vsNR clusters in the data set.

### Blinking analysis and thresholding

The PL intensity distribution of an individual vsNR contains temporal intermittency (blinking) pattern[58] and it is composed of so-called "on" states and "dark" states. During "on" states the particle PL is bright, whereas during "dark" states it is mostly background noise fluorescence. The characterization of blinking periods is particularly hard, with heavy-tail power-law distributions.[68] In our experiments this translated to a finite fraction of the recording with no signal coming from the vsNR, see an example in Fig. S3, or even frames displaying an intermediate state – a mixture of "on" and "dark" states that last less than the acquisition rate, 10 ms. To improve the quality of vsNR signal, we developed a routine to identify and excise the "dark" state frames from the recording.

To classify the two states, we calculated for each ROI the distribution of PL intensity after background subtraction. In Fig. S3b, this distribution is shown for a time trace from S3a.

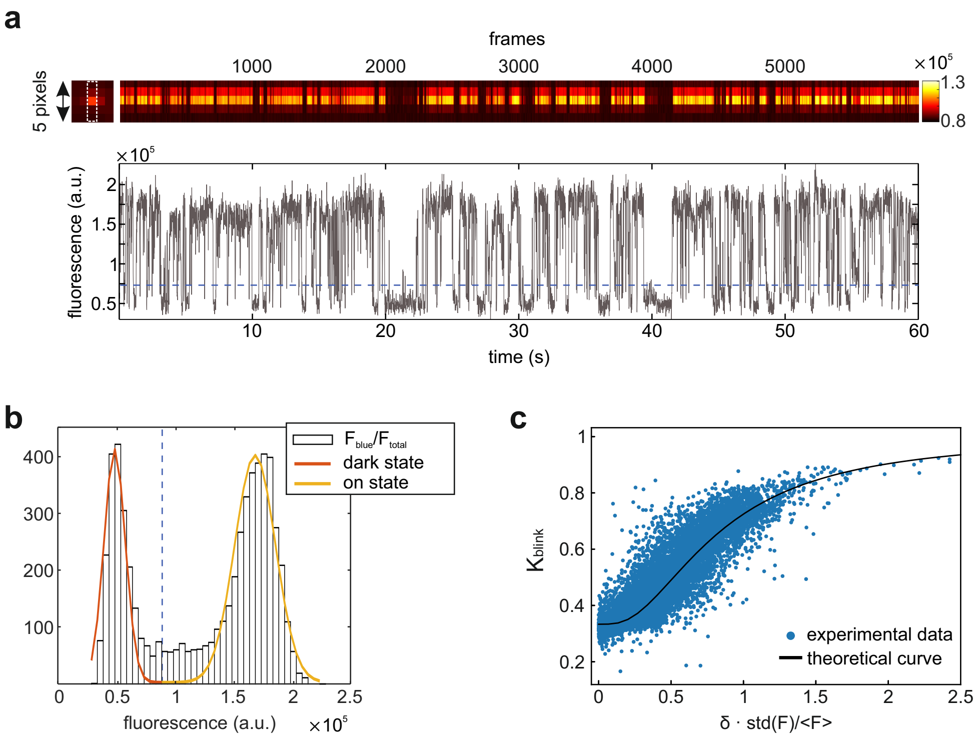


Figure S3: (a) An example of a blinking vsNR time trace. Upper panel shows color-coded intensity profile of an ROI over 60 s (6000 frames). The intensity was collected from 5 pixels highlighted by dashed white rectangle on the ROI image on the left. Bottom panel depicts time trace of the same vsNR after full processing including background subtraction and summation of $F_{green}$ and $F_{red}$ channels. Blue dashed line corresponds to the blinking threshold. (b) Distribution of vsNR intensities collected during 60 s (6000 frames), the same recording as in (a). (c) Blinking coefficient (eq. 1) for a selection of 17500 local maxima as a function of the fitted $\delta$ in the bimodal Gaussian distribution, eq. 6, in units of the average fluorescence. The solid line represents the theoretical blinking coefficient for the ideal case of two Gaussian distributions with $p = 1/2$ and $\sigma_1 = \sigma_2 = 1$ as a function of $A\delta$, with $A \approx 0.167$.

We fitted the full distribution, after regularisation[1], to a sum of two Gaussian distributions,

$$f(x) = \frac{p e^{-\frac{(x-\mu_0)^2}{2\sigma_1^2}}}{\sqrt{2\pi\sigma_1^2}} + \frac{(1-p) e^{-\frac{(x-(\mu_0+\delta))^2}{2\sigma_2^2}}}{\sqrt{2\pi\sigma_2^2}} \quad , \tag{6}$$

where $\mu_0$, $\delta$, $\sigma_1$, $\sigma_2$ and $p$ are free parameters. The part of the distribution corresponding to the "dark" state is centered at $\mu_0$ and its width is given by $\sigma_1$. $\mu_0 + \delta$ and $\sigma_2$ are the

[1]To regularise data for fitting to distributions we subtracted the average and divided by the standard deviation.

parameters of the "on" state. We defined a threshold to excise the "dark" state frames as 2 times $\sigma_1$ from the center of the "dark" state $\mu_0$. In a few cases, the "dark" state distribution was too broad, due to a higher probability of mixed states. To avoid discarding these cases, we considered that if the threshold is above $\mu_0 + \delta$, it would be replaced by $\mu_0 + \delta/2$, the middle point in between the two peaks of the distribution. The fitted function is shown in Fig. S3b as a solid line (in red the "dark" state part and in yellow the "on" state).

Using eq. 6 as the distribution, we computed the theoretical blinking coefficient, eq. 1, for an ideal trace with two equally probable on and off states ($p = 1/2$ and $\sigma_1 = \sigma_2 = 1$) and compared it with the extracted blinking coefficients for randomly selected local maxima, Fig. S3c. Blinking coefficients above 0.45 correspond to interstates distance larger than $\delta > 2.5\ \sigma_1$.

### Spectral separation between vsNR and BeRST channels

To verify efficient separation between vsNR and BeRST signals we performed a control experiment when we loaded neurons only with BeRST (Fig. S4a) and recorded vsNR channel (Fig. S4b) using our normal protocol. BeRST fluorescence closely followed membrane voltage modulation (Fig. S4c), whereas the signal collected inside the BeRST mask on the vsNR channel showed no correlation with the modulation (Fig. S4d). We have also performed Fast Fourier transform of the signal collected on the vsNR channel and we found no band corresponding to the frequency of voltage modulation (25 Hz, Fig. S4e). Thus, we concluded that the separation between the two channels was efficient and there was no leakage of BeRST fluorescence into vsNR channel.

### FT score and bootstrap

To quantify uncertainty in our data we proceeded to generate synthetic traces that reproduced the statistical features of the traces shown in Fig. 3c. We modeled the trace, $S$, as a

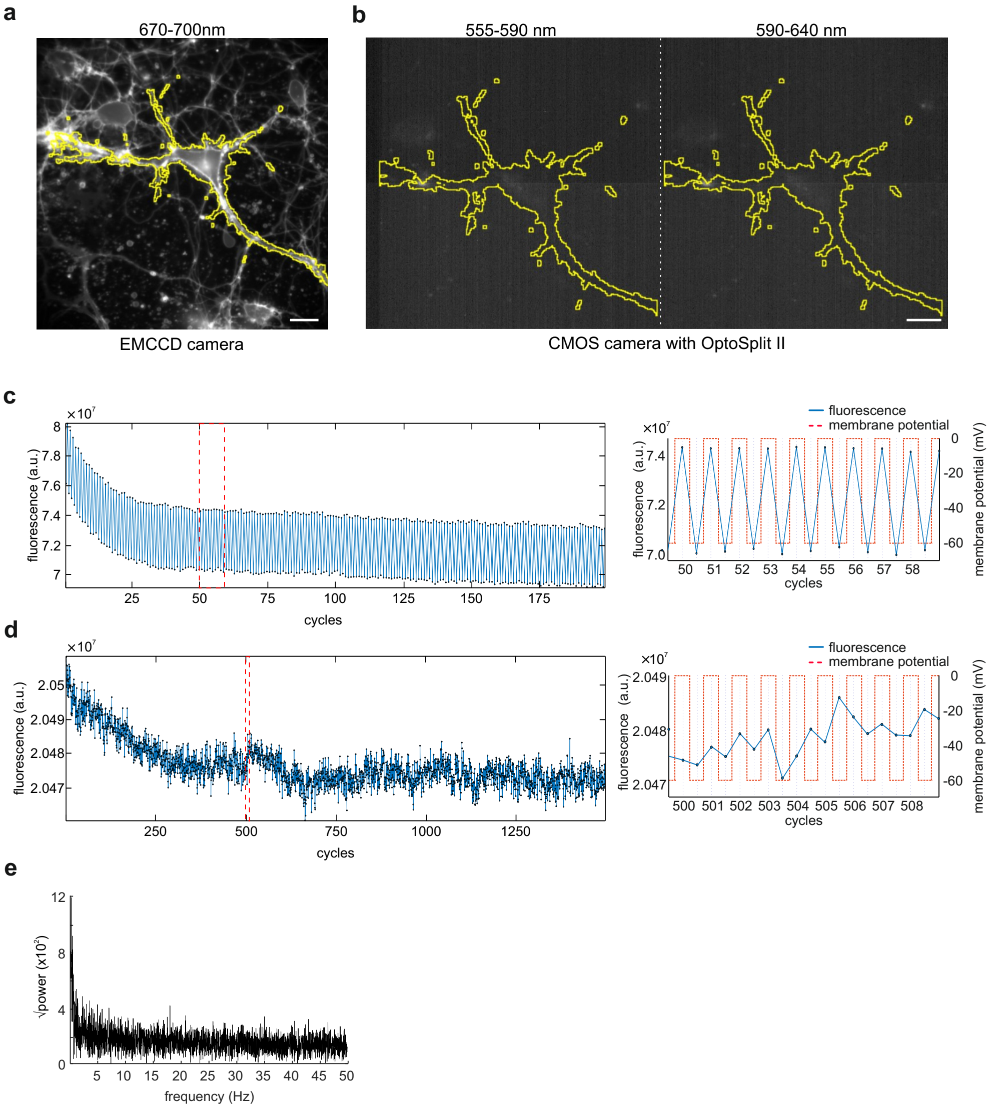


Figure S4: Spectral separation of vsNR and BeRST channels. (a) BeRST-loaded neuron with the BeRST mask overlay (yellow line). (b) Split emission image of the same field as in (a). Since vsNRs were not loaded in this experiment there is no fluorescent signal in this channel. (c) Fluorescent time trace of the BeRST signal inside the mask. The left panel depicts the entire recording, whereas the right panel shows an enlarged part of the trace designated with the red dashed rectangle. BeRST fluorescence (blue line) closely followed membrane voltage modulation (dashed red line). (d) Time trace of the signal collected inside the BeRST mask on the vsNR channel. The trace corresponds to the sum of the two vsNR split emission channels. The left panel depicts the entire recording, whereas the right panel shows an enlarged part of the trace designated with the red dashed rectangle. The absence of the leakage of BeRST signal into vsNR channel is evident from the absence of the correlation between fluorescence in vsNR channel and voltage modulation. (e) Fast Fourier transform of the signal depicted in (d). Note the absence of the 25 Hz band corresponding to the frequency of membrane voltage modulation (see fig. 6b for comparison).

Gaussian process centered around a mean, $\mu$, and with Gaussian white noise of width $\sigma$.

Given the heterogeneity of the NR responses, using a single pair of values $\mu$ and $\sigma$ (say ensemble average and standard deviation) did not reflect the variability and properties of the signals we have analyzed. Instead, we considered bootstrap methods to keep the correlated statistics of the traces. First, we extracted the average and standard deviation of the selected traces, once the blinking periods had been excised, and constructed a set of pair values $\mathcal{D} = \{(\mu_i, \sigma_i)\}_{i=1,\ldots,N}$. Then, we resampled the distribution of parameters extracted from the data. To do that, for each new synthetic trace we generated a random index, $j$, with its associate pair $(\mu_j, \sigma_j)$. We added to the selected values small Gaussian noise, to smooth out the distribution of parameters. The noise level was set such that the distributions remained similar to the original ones. Particularly, 10% noise was enough to do it without perturbing correlations. The duration of the traces was also set by resampling the distribution of length of the traces stitched after blinking periods excision. This procedure produced synthetic traces similar to the signal traces, with the same overall statistics.

Subsequently, we used the generated synthetic traces to set the definition of the FT score: the ratio between the value of the peak at the target frequency, 25 Hz, and the standard deviation of the surrounding frequency bandwidth. To set the bandwidth, we generated $10^5$ synthetic traces for each bandwidth and study the percentage of traces with (bandwidth-dependent) FT score above 5 (data not shown). This curve flattened out at around 6 Hz. We used 10 Hz as bandwidth for the definition of the FT score. For this particular value, we studied the distribution of the FT score, by generating $10^6$ synthetic traces. The distribution is shown in Fig. S5a, and in semi-logarithmic scale in Fig. S5b. More precisely, the integral of this curve, namely the p-value, shows the probability of having a trace without signal and with FT score above a threshold, Fig. S5c.

For FT-score value of 5, p-value was below 0.001 (roughly $p_{FT} \approx 8 \cdot 10^{-4}$), i.e., the probability of finding a spurious trace with FT score 5 was below 0.1%. We found 8 vsNRs over 1242 with FT score above 5, the probability that they are all spurious can be calculated

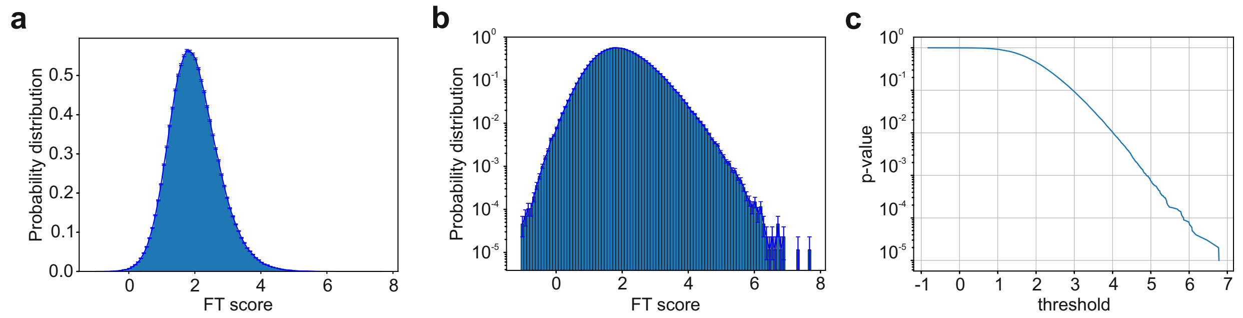


Figure S5: FT score of bootstrapped traces. (a) Probability distribution of FT score values estimated from $10^6$ bootstrapped synthetic traces, in linear scale. (b) the same in semi-logarithmic scale. (c) The $p$–value of the probability distribution shown in (a) and (b). This is the probability that a given trace will be above the threshold, represented in the x-axis.

from Bayes' theorem. If we estimate the probability of finding signal in the selected traces of $p_s \sim 0.001$, then the probability that a given trace with FT score above 5 is spurious was roughly $(1 + \frac{p_s}{p_{FT}(1-ps)})^{-1} \approx 0.44$. Since they are independent, the probability that the 8 selected traces do not contain any signal is 0.0015.

### Burst intervals

To find periods of voltage sensitivity in signal of vsNR with FT score greater than 5, we used the following burst-search algorithm. We summed the green ($F_{\text{green}}$) and the red ($F_{\text{red}}$) components of the raw signal (Fig. S6a) to produce $F_{\text{total}}$ (F(t) in eq. 2). $F_{\text{total}}$ was used to find blinking threshold as described in SI section "Blinking analysis and thresholding" (Fig. S6b). Blinking periods were excised and any cycle of the stimulation square wave that contained at least one blinking frame was removed. The rest of the trace was stitched, forming the sequence of "on" frames that preserved the cycles of the stimulation wave. We refer to this trace as an "aligned trace". Identical frames were stitched for $F_{\text{total}}$ and Rt traces (Rt(t) in eq. 2, Fig. S6c). Next, we calculated ΔF/F and ΔRt using aligned $F_{\text{total}}$ and Rt traces by averaging frames with identical voltage in each cycle and calculating the difference between -60 mV and 0 mV steps. ΔF was additionally normalized to $F_{\text{-60mV}}$ in each cycle (eq. 3). From ΔF/F and ΔRt we calculated a running mean with a window of

10 cycles of the stimulation square wave (red line in Fig. S6d and e). At the next step, we identified positive (running mean above 0) and negative (running mean below 0) intervals and calculated the burst score as the integral of the running mean for these intervals. Finally, we selected most prominent intervals by using a threshold calculated individually for each particle based on the control trace burst score values distribution (dashed red rectangles in Fig. S6c, d, and e). Since burst-search algorithm relies on a thresholding of the running mean, it occasionally produces false-positive intervals, and should be used only in combination with the Fourier Transform analysis to identify the putative time positions where the 25 Hz component dominates.

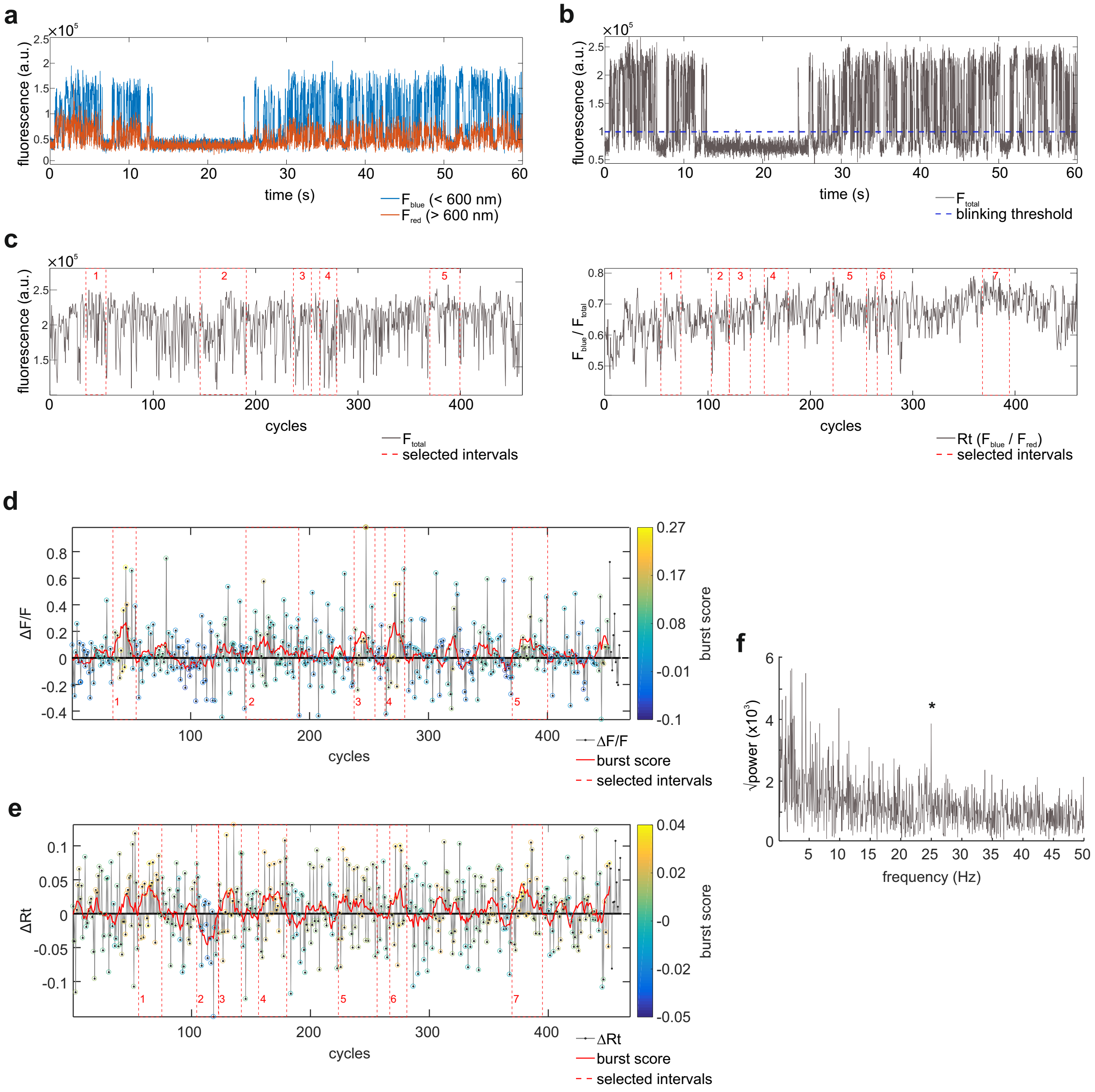


Figure S6: Selection of responsive intervals using the burst score. All traces in this figure belong to the same vsNR as the one shown in Fig. 7. (a) Time trace of the fluorescent signal of the vsNR in reflected ($F_{green}$) and transmitted ($F_{red}$) channels produced by a dichroic mirror with 600 nm edge. (b) Sum of reflected and transmitted channels ($F_{total}$) with blinking threshold (dashed blue line). (c) $F_{total}$ (left panel) and ratio (Rt, $F_{green}/F_{total}$, right panel) time traces. To produce these traces blinking periods were excised and cycles of the stimulation square wave were aligned. (d) $\Delta F/F$ time trace. Burst score (running mean of $\Delta F/F$ with window of 10 cycles) is depicted as a red line. Intervals selected based on the burst score magnitude are highlighted with dashed red rectangles. These intervals are also highlighted in (c), left panel, and shown magnified in Fig. 6c. (e) The same as (d) but for the $\Delta$Rt time trace. $\Delta$Rt intervals are also highlighted in (c), right panel, and shown magnified in Fig. 6d. (f) Fast Fourier transform of the $F_{green}$ signal. 25 Hz band corresponding to the frequency of membrane voltage modulation is highlighted with an asterisk.

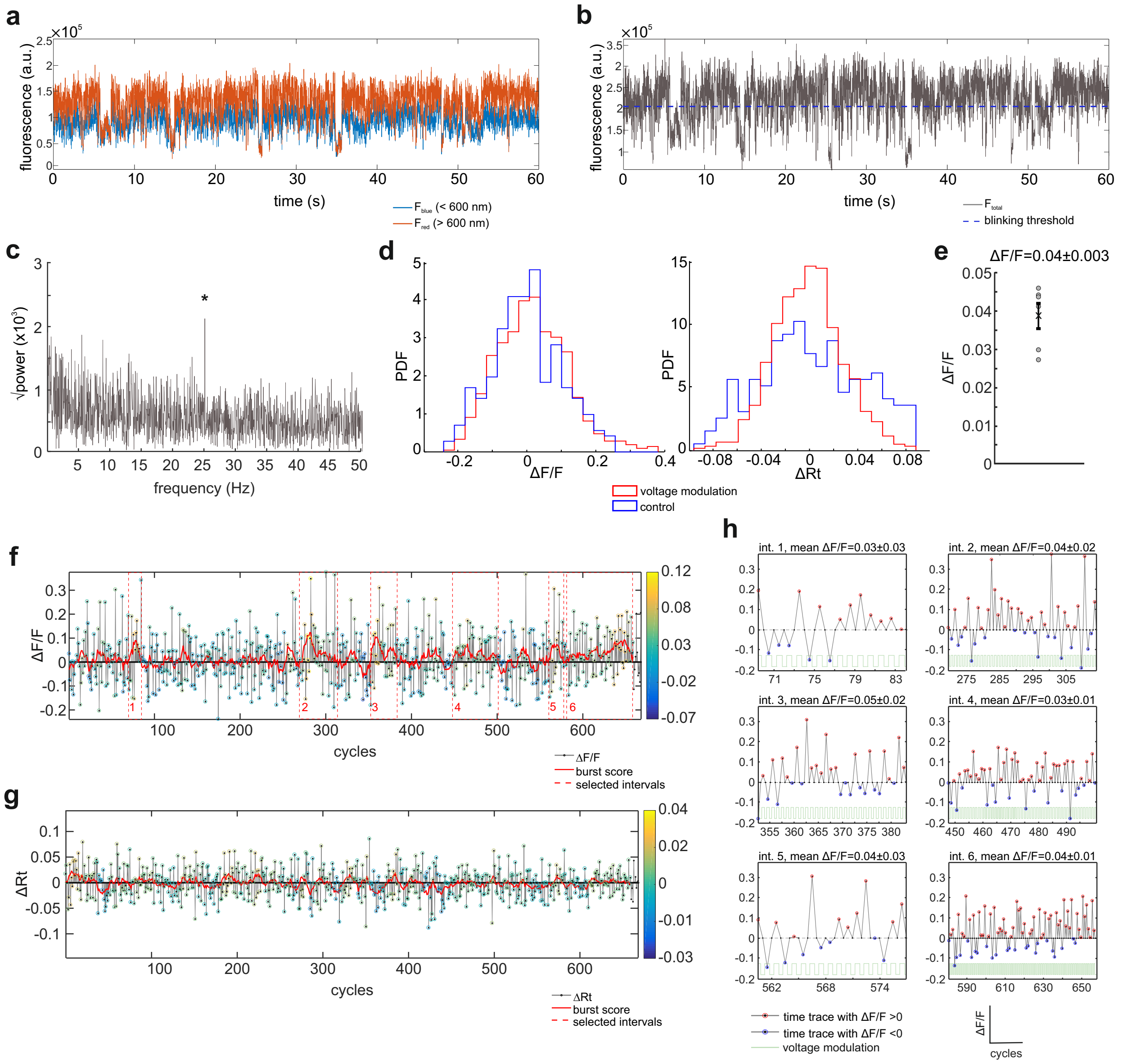

Figure S7: All traces in this figure belong to the same vsNR, different from the one shown in Fig. 7 and Fig. S6. (a) Time trace of the fluorescent signal of the vsNR in reflected ($F_{green}$) and transmitted ($F_{red}$) channels. (b) Sum of reflected and transmitted channels ($F_{total}$) with blinking threshold (dashed blue line). (c) Fast Fourier transform of the $F_{green}$ signal. 25 Hz band corresponding to the frequency of membrane voltage modulation is highlighted with an asterisk. (d) Distribution of $\Delta F/F$ (left panel) and $\Delta Rt$ (right panel) of the vsNR with (red) and without (blue) voltage modulation. (e) Mean $\Delta F/F$ of each intervals selected from the trace with voltage modulation. The mean of all selected intervals is indicated with the cross, error bars depict STE. No $\Delta Rt$ intervals were found for this vsNR (see (g)). (f) $\Delta F/F$ time trace. Burst score is depicted as a red line. Selected intervals are highlighted with dashed red rectangles. These intervals are also shown magnified in (h). (e) The same as (d) but for the $\Delta Rt$ time trace. No $\Delta Rt$ intervals were found for this vsNR. (h) All selected intervals of $\Delta F/F$ time trace.
